# A conservative moving-boundary multiphysics framework reveals nonlinear coupling between osmotic flow and actin-driven cell migration

**DOI:** 10.64898/2026.10.09.757961

**Authors:** Han Zhou, Lingxing Yao, Yoichiro Mori, Yizeng Li

## Abstract

Cell migration couples membrane transport, intracellular fluid motion, and actin remodeling on a boundary that deforms with the cell. Quantitative interpretation of this coupling requires accurate one-sided membrane solute transport and conservative transfer between the G-actin and F-actin pools, because these fields determine the osmotic pressure and polymerization flux that enter the force balance. We develop a semi-implicit immersed-boundary Stokes–Brinkman solver coupled to a sharp interface Cartesian grid advection–diffusion solver that enforces Robin conditions on the evolving membrane and advances F-actin/G-actin exchange through paired interface fluxes on the same accepted geometry. The framework resolves membrane mechanics, osmotic water permeation, solute transport, actin turnover, substrate drag, and intracellular advection in one calculation. Dedicated actin-driven and pump-induced tests show excellent conservation of total actin and total solute, respectively, under sustained interface motion and transport. The actin calculations recover biphasic dependencies on both depolymerization and substrate coupling, while pump-induced migration generates a rearward cell-relative cytosolic flow. In the fully coupled system, cell deformation generated by actin polymerization interacts with active solute pumps to produce further changes in morphology and cell area. Cell expansion reflects accumulated net water entry at fixed hydraulic permeability. Matched controls show that the coupled actin–osmotic system travels farther than the baseline-corrected sum of the isolated actin-driven and pump-induced responses. Positive excess displacement accompanies a longer span of cell-relative retrograde F-actin motion and a greater magnitude of total rearward substrate force. These changes arise from the coupled hydraulic and mechanical response of the deforming cell.

## 1 Introduction

Cell migration underlies embryonic morphogenesis, wound repair, immune surveillance, angiogenesis, and cancer invasion and metastasis [1–4]. A migrating cell must coordinate polarization, actin-generated force, adhesion turnover, membrane deformation, intracellular transport, and volume regulation as its shape changes [2, 4–6]. Actin polymerization against the membrane and myosin-driven network contraction generate cytoskeletal stresses [5]. Mechanical coupling among the network, membrane, and cytosol transmits these stresses and drives intracellular flow [6, 7]. A quantitative model of these interactions must resolve the moving cell geometry while maintaining conservation of the transported species as mechanics and transport evolve together [8–11].

Actin network organization and force transmission through the membrane and adhesions regulate cell shape and migration [5, 12–15]. Globular actin (G-actin) forms the soluble monomer pool, whereas filamentous actin (F-actin) forms dynamic networks that generate and transmit mechanical stress [12–14]. Polymerization consumes G-actin at sites of filament assembly and depolymerization returns monomers to the soluble pool, so sustained protrusion depends on both actin turnover and transport of monomers to regions of active assembly [12, 13, 16, 17]. Directed intracellular transport can supplement diffusion in the delivery of soluble G-actin and other proteins [16–19]. Water and solute transport provide a second source of cell motion through ion transporters, channels, aquaporins, and local volume regulation [4, 20]. NHE1 (Na^+^/H^+^ exchanger 1) is a plasma-membrane ion exchanger, and SWELL1 (LRRC8A) is an essential subunit of the volume-regulated anion channel [20, 21]. Their front and rear enrichment, respectively, supports local swelling and shrinkage in confined breast cancer cells; optogenetic redistribution of SWELL1 changes migration direction and efficiency [22]. Polarized solute uptake and release establish osmotic pressure differences that drive water entry at the front and exit at the rear [20, 22]. Our model represents this polarized transport with a neutral solute. Intracellular fluid flow has also been measured in migrating cells and can advect soluble material, which creates a physical route from membrane transport to intracellular concentration fields [6, 7, 19].

Our earlier theoretical studies established several components of this physics separately. Two-phase models analyzed actin-driven and pump-induced migration as complementary mechanisms and quantified how their energetic efficiency depends on the hydraulic environment [23, 24]. Pump-induced migration denotes the same mechanism termed water-driven migration in our earlier work: active solute pumping creates an osmotic imbalance that drives transmembrane water flux. Subsequent actin models introduced G-actin transport, depolymerization, and two-dimensional morphology, but we did not couple these dynamics to a cytosolic flow field generated by polarized osmotic transport [25–27]. The unresolved problem is the dynamical coupling among osmotic water permeation, cell deformation, actin dynamics, and substrate force transmission. Resolving this feedback requires solute, cytosol, F-actin, G-actin, and a deforming membrane to evolve as one coupled system. Membrane geometry enters the normal mechanical terms of the water-permeation law, while substrate drag transmits the force generated across the evolving actin distribution.

A direct calculation of this transport–mechanics feedback requires partial differential equations on a domain whose boundary moves as part of the solution [11, 28, 29]. Body-fitted arbitrary Lagrangian– Eulerian formulations impose interface conditions on a conforming mesh and are well established for fluid–structure interaction, while immersed formulations allow large interface motion on a fixed Eulerian grid [28–31]. Accurate preservation of enclosed volume under immersed Lagrangian–Eulerian coupling has motivated divergence-free interpolation and adjoint force-spreading strategies [9, 10]. For the present problem, however, geometric accuracy alone is insufficient. The membrane is simultaneously a force-bearing boundary, a semipermeable interface, and a site of actin conversion, so the mechanical and solute boundary conditions must remain registered to the same evolving geometry.

Interfacial transport imposes the more restrictive requirement. A permeable moving membrane supports one-sided concentrations, normal derivatives, and Robin-type flux conditions that determine the osmotic pressure and hence the fluid motion [11, 32–34]. Immersed mass-transfer, embedded-boundary, cut-cell, immersed-interface, and unfitted finite-element formulations provide different routes to resolve transport on deforming or irregular domains [11, 32–38]. Recent Cartesian formulations extend sharp Robin transport to evolving domains without body-fitted remeshing [39]. In the cell-migration problem studied here, the one-sided solute fluxes determine the concentration jump that drives water permeation, while the F-actin/G-actin exchange determines the monomer supply available for membrane polymerization. The numerical treatment must preserve both boundary flux accuracy and actin conservation on the same evolving interface.

Material conservation provides an integrated test of these coupled numerical operations. Total actin is conserved at the continuum level while F-actin and G-actin move at different velocities, interconvert in the bulk, and exchange through paired polymerization fluxes on a deforming membrane. Total solute is likewise conserved while one-sided Robin fluxes transfer material across the membrane and osmotic water permeation changes the intracellular geometry. We quantify the relative conservation error in total actin and total solute in addition to conventional spatial and temporal accuracy studies [9–11, 33, 34].

In this work, we develop a moving-boundary formulation in which fluid mechanics, membrane deformation, solute transport, osmosis, F-actin/G-actin interconversion, substrate drag, and intracellular advection share one evolving membrane geometry. The numerical architecture is organized around the corresponding biophysical requirements: semipermeable membrane mechanics, sharp transmembrane transport, and paired F-actin/G-actin exchange on one accepted moving interface (Fig. 1A). The shared interface allows solute transport, normal membrane forces, and actin dynamics to determine water permeation and cell deformation together. We establish conservation independently in the actin and osmotic subsystems over the time scales used for migration before activating both mechanisms. The coupled calculation resolves the resulting changes in cell area, actin motion, and substrate force transmission.

**Figure 1:**
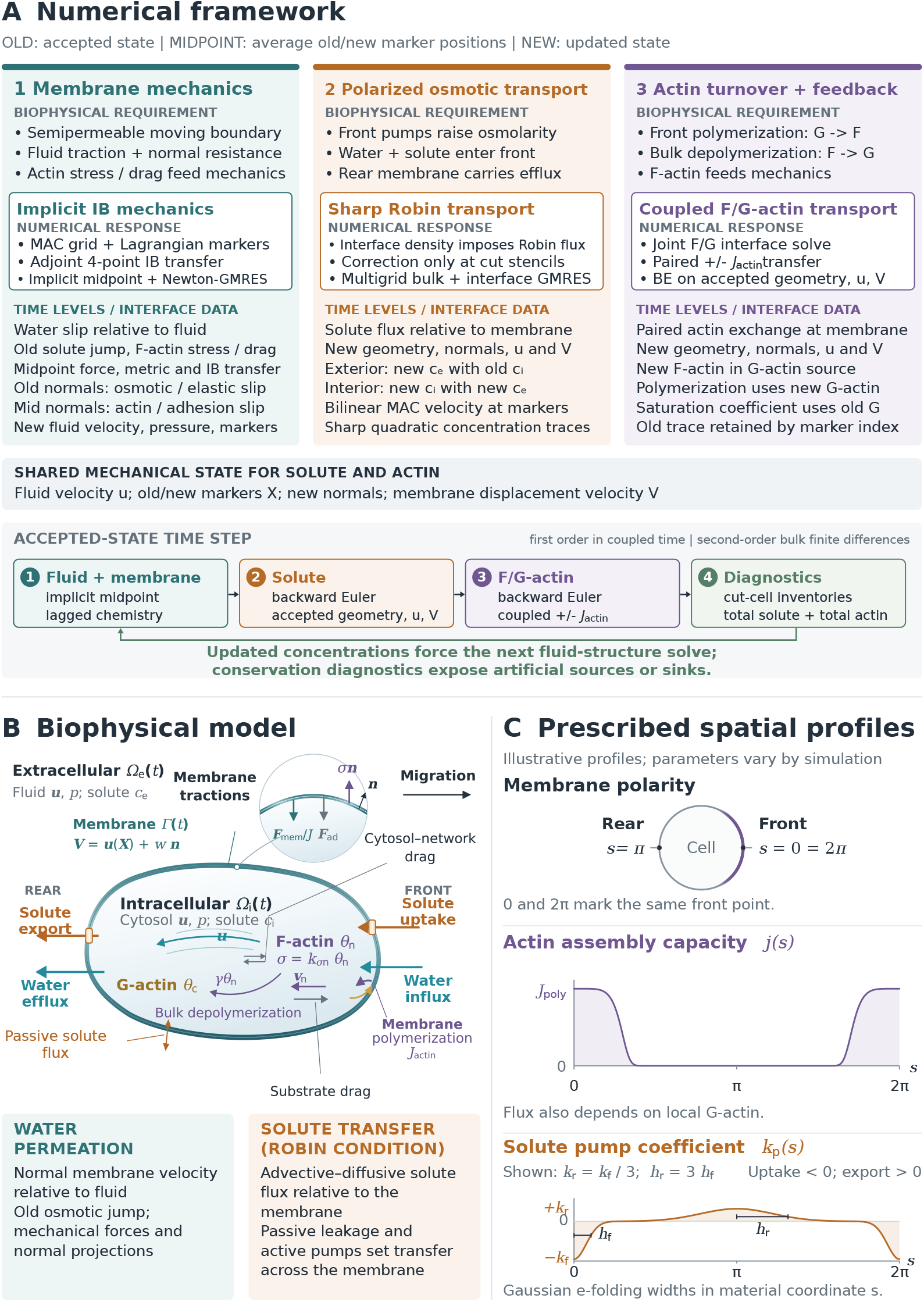
Numerical framework, biophysical model, and prescribed profiles. **(A)** Sequential fluid– membrane, solute, and coupled actin updates. Mechanics uses lagged concentrations and midpoint membrane positions; transport uses the accepted mechanical state. Old and new denote accepted and updated states. **(B)** Cellular domains, actin exchange, and mechanical interactions. Water permeation determines membrane motion relative to fluid; the Robin condition prescribes solute transfer. **(C)** Membrane polarity, assembly capacity *j*(*s*), and signed Gaussian pump profiles. The front is at *s* = 0 or 2*π* and the rear at *s* = *π*. Uptake has *k*_p_ *<* 0 and export *k*_p_ *>* 0; the broad-rear profile uses *k*_r_ = *k*_f_*/*3 and *h*_r_ = 3*h*_f_.

## 2 Methods

We consider a two-dimensional cell as a closed deformable membrane that separates the intracellular domain Ω_*i*_(*t*) from the extracellular domain Ω_*e*_(*t*). We denote the membrane by Ω(*t*) and the fixed fluid domain by Ω = Ω_*i*_(*t*) *∪* Ω(*t*) *∪* Ω_*e*_(*t*). The membrane is parameterized by the material map ***X***(*s, t*), with outward unit normal ***n***, stretch *J* = |*∂*_*s*_***X***|, and membrane velocity ***V*** = *∂*_*t*_***X***. The fluid velocity and pressure are ***u*** and *p*. The intracellular actin system contains F-actin concentration *θ*_*n*_ and G-actin concentration *θ*_*c*_, with initial concentrations *F*_0_ and *G*_0_, respectively; the neutral solute concentration is *c*_*i*_ inside and *c*_*e*_ outside. The same membrane geometry carries mechanical traction, osmotic and solute fluxes, and actin exchange (Fig. 1B). Solute and actin concentrations enter the mechanical balance and water-permeation law, while membrane motion sets the relative normal transport at the interface.

### 2.1 Coupled continuum model

The F-actin network is treated as a porous mechanical phase that exchanges drag with the cytosol and with the substrate. The network velocity is ***v***_*n*_, *σ* is the F-actin network stress, and *k*_*σn*_ is the coefficient of F-actin pressure in the linear constitutive law [25, 26]. The cytosol exerts interphase drag *ηθ*_*n*_(***u*** *−* ***v***_*n*_), and focal adhesions transmit substrate drag *−η*_st_*θ*_*n*_***v***_*n*_, which opposes network motion. Here *η* and *η*_st_ are drag coefficients per unit F-actin concentration. The network force balance is [23, 24, 27]

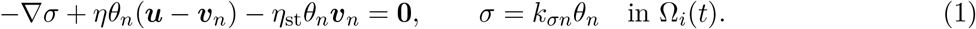

Solving the network force balance for the F-actin flux gives

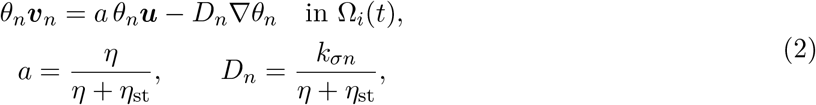

where *a* is the fraction of cytosol advection inherited by the F-actin network and *D*_*n*_ is its effective diffusivity. At fixed *η* and *k*_*σn*_, increasing *η*_st_ lowers both *a* and *D*_*n*_, reducing cytosol-driven and stress-driven network fluxes for fixed velocity and concentration fields.

At the cellular length and velocity scales considered here, inertia is neglected. The fluid satisfies a Stokes–Brinkman balance in which *µ* is the dynamic viscosity, *χ*_*i*_ is the indicator of the intracellular domain and confines actin-dependent drag and forcing to the interior, and ***f***_Ω_ is the force density spread from the membrane to the fluid grid. The governing equations are

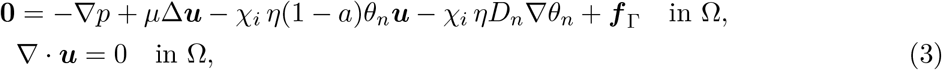

where ***x*** is the Eulerian position and *δ* is the Dirac distribution. The membrane elastic force ***F***_mem_, actin stress at the membrane, and membrane–substrate normal resistance ***F***_ad_ are

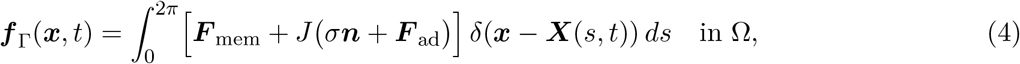

where *k*_m_ is the membrane elastic modulus in the linear membrane tension law *T*_m_ = *k*_m_*J*, ***τ*** is the unit tangent, and *k*_ad_ is the membrane normal-resistance coefficient. The F-actin network stress at the membrane uses the interior trace of *θ*_*n*_. The force ***F***_mem_ is defined per unit material coordinate, and ***F***_mem_*/J* gives force per unit current membrane length. The membrane and adhesion forces follow the two-phase migration and two-dimensional actin formulations developed previously [24, 27].

The membrane is permeable to water. Its velocity equals the local cytosol velocity plus a normal permeation velocity *w*, defined by *w* = (***V*** *−* ***u***(***X***)) *·* ***n***. Positive *w* denotes water entry, since the outward fluid flux relative to the membrane is *−w*. The hydraulic permeability is *k*_w_, *R* is the gas constant, *T* is absolute temperature, and [*c*] = *c*_*i*_ *− c*_*e*_ is the interior-minus-exterior solute concentration jump. The membrane kinematics are

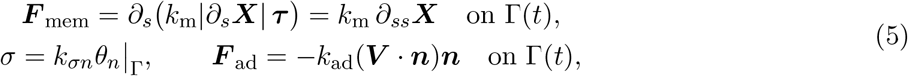

The linear hydraulic law describes passive permeation driven by the ideal-solution osmotic pressure jump *RT* [*c*] and normal mechanical traction from actin stress, membrane elasticity, and membrane resistance [11, 24, 25].

The neutral solute diffuses and is advected by the fluid on each side of the membrane. For *ℓ ∈ { i, e, c*_*ℓ*_*}* is the one-sided solute concentration and *D*_s_ its diffusivity. The passive permeability *k*_sol_ sets the signed outward flux *k*_sol_ [*c*], a linear approximation to chemical-potential-driven permeation near *c*_*i*_ = *c*_*e*_ [11, 25]. The active coefficient *k*_p_(*s*) is prescribed in membrane material coordinates; both coefficients have units of velocity. Positive *k*_p_ denotes export and negative *k*_p_ denotes uptake. The donor trace is *H* = *c*_*i*_ where *k*_p_ *≥* 0 and *H* = *c*_*e*_ where *k*_p_ *<* 0, so the active flux *k*_p_*H* vanishes upon donor depletion. The transport law is

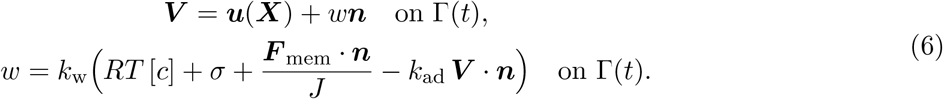

so the same interfacial transfer appears in the two one-sided boundary conditions. Using the same transfer on both sides conserves the total amount of solute at the continuum level while allowing active redistribution between the intracellular and extracellular domains [11, 24].

F-actin and G-actin occupy the intracellular domain and exchange material through bulk depolymerization and membrane polymerization. The depolymerization rate is *γ*, the G-actin diffusivity is *D*_*c*_, *j*(*s*) is the prescribed leading-edge polymerization-capacity profile, and *θ*_*c,c*_ is the G-actin concentration at which the membrane polymerization flux reaches half maximum. The amplitude of *j*(*s*) is *J*_poly_. The coupled actin equations are [27]

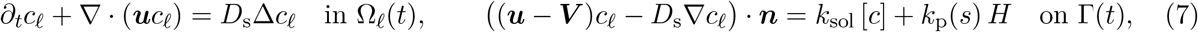

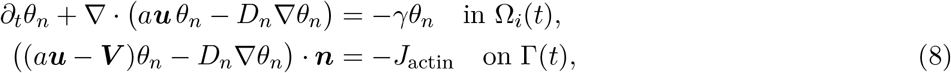

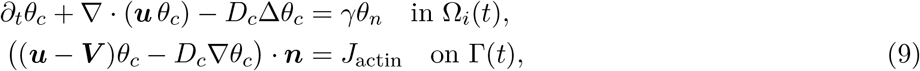

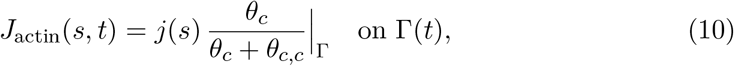

The equal and opposite polymerization fluxes in the two boundary conditions convert G-actin into F-actin at the leading membrane while conserving total actin. Bulk depolymerization converts F-actin into G-actin at the rate *γθ*_*n*_; *γ* = 0 sets both the F-actin sink and the G-actin source to zero. The F-actin flux combines advection by *a****u*** with effective diffusivity *D*_*n*_; G-actin is advected by ***u*** and diffuses with coefficient *D*_*c*_. F-actin enters the fluid problem through drag and stress, which closes the mechanical feedback. The osmotic and actin subsystems couple directly through the shared cytosolic velocity, concentration fields, and moving boundary. Fluid velocity and pressure are determined throughout Ω, solute concentrations on their respective sides of the membrane, and both actin concentrations in Ω_*i*_(*t*); membrane position, forces, and fluxes are determined on τ(*t*).

### 2.2 Moving-boundary numerical method

The numerical method mirrors this biophysical structure and uses one geometrical representation of the membrane for every subsystem (Fig. 1A). Fluid velocity is stored on a staggered marker-and-cell (MAC) Cartesian grid, pressure and concentrations are cell centered, and the membrane is represented by Lagrangian markers that move through the fixed Eulerian mesh. Membrane forces are spread to the fluid and fluid velocities are interpolated to the markers with the same four-point regularized delta kernel, which gives an adjoint pair of transfer operators in the immersed-boundary discretization [30]. Membrane elasticity, normal resistance, and water permeation are treated within a semi-implicit fluid–structure update, following the stability strategy of implicit immersed-boundary methods and the osmotic moving-membrane formulation used in earlier work [11, 40]. The mechanical update advances the geometry on which all subsequent transport conditions are imposed.

Solute and actin transport require sharp one-sided boundary conditions that differ from the regularized force transfer used by the fluid solver. We reformulate each moving-domain transport problem on the fixed Cartesian box with an interface density that enforces the Robin condition. A local quadratic correction modifies only finite-difference stencils cut by the membrane; away from the interface, the standard Cartesian advection–diffusion operator is unchanged. This sharp treatment is important here because the osmotic driving is set by the one-sided concentration jump itself; numerical smearing across the membrane would directly alter the water flux that generates intracellular flow. The fixed-grid formulation avoids body-fitted remeshing, and confining the correction to intersected stencils preserves the Cartesian bulk operator used by geometric multigrid and the matrix-free interface solve [39]. F-actin and G-actin interface densities are solved as one coupled vector so that their equal-and-opposite polymerization fluxes are applied at the same membrane state. The conversion of G-actin into F-actin is numerically paired at the interface instead of being assembled from independently evaluated transfers.

Each time step advances the fluid–membrane system first, followed by solute transport and then the coupled actin transport. The fluid–membrane subproblem uses a semi-implicit midpoint scheme: the membrane configuration and elastic force are determined implicitly, with the solute jump and F-actin concentration fixed at the previous time level. Backward Euler transport and this lagged feedback make the coupled scheme first order in time. The bulk finite-difference operators are second order in space. The same accepted membrane geometry, fluid velocity, and membrane velocity are passed to the solute and actin solves, and the updated solute fields then force the next fluid–structure step. Discrete operators, solver tolerances, spatial and temporal resolutions, and the translating-window algorithm are specified in Supplementary Sections S2–S4; physical parameters and units are listed in Section S1.

We track total actin and total solute because their conservation is directly coupled to membrane polymerization and osmotic pressure in the model. In the continuum equations, both totals are conserved in the absence of external sources. Their discrete conservation depends jointly on the accepted membrane geometry, one-sided interface reconstruction, cut-cell integration, advection, diffusion, species exchange, and transmembrane fluxes. Total actin is the intracellular integral of *θ*_*n*_ + *θ*_*c*_, and total solute is the sum of the intracellular and extracellular solute integrals over the computational domain. Fully interior grid cells contribute their full area; cells cut by the membrane use geometrical cell fractions and one-sided reconstructed concentrations. We define the relative conservation error from the change in these total amounts from their initial values. This provides a direct quantitative measure of conservation accuracy for the complete moving-boundary transport update.

### 2.3 Simulation design and quantitative diagnostics

All calculations begin from a circular cell with uniform solute concentration on the two sides of the membrane. Actin polymerization is localized near the leading edge in membrane material coordinates. Active solute pumping is prescribed by Gaussian profiles at the front and rear of the cell. Their positive magnitudes are denoted *k*_f_ for front uptake and *k*_r_ for rear export, with material-coordinate widths *h*_f_ and *h*_r_; the signed coefficient *k*_p_(*s*) in Eq. (7) carries the corresponding transport direction (Fig. 1C). All reported migration calculations begin with *F*_0_ = *G*_0_ = 0.1 mM. The standalone pump-induced calculations set *J*_poly_ = 0 to suppress membrane polymerization and *γ* = 0 to eliminate bulk F-actin-to-G-actin conversion. The concentrations continue to evolve through advection and diffusion, and each actin pool is separately conserved at the continuum level. Complete parameter sets are listed in Supplementary Tables S1 and S2.

Cell position is measured by the area centroid. For actin-driven migration, velocity is calculated from the centroid displacement over the stated analysis interval. Pump-induced and coupled calculations report laboratory-frame displacement relative to the stationary substrate; when a translating computational window is used, its accumulated translation is restored. Cell-relative velocities subtract the laboratory-frame centroid velocity; the fitting procedure for the coupled centerline profiles is specified in Supplementary Section S4. Actin polarity is defined as 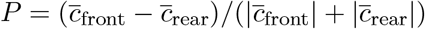, where the bars denote arithmetic means of intracellular grid samples on the front and rear sides of the instantaneous area centroid, and *c* = *θ*_*n*_ or *θ*_*c*_ for F-actin or G-actin. Positive *P* denotes front enrichment and negative *P* rear enrichment.

Actin polymerization and active solute pumping drive migration through different physical processes in the uncoupled limits of the model. Polymerization supplies F-actin at the leading membrane; the resulting network stress and substrate drag convert actin motion into cell translation. Polarized active solute pumps move solute across the membrane, which creates asymmetric osmotic pressure and drives transmembrane water flux and intracellular flow.

The matched simulations combine two settings for actin turnover with two settings for active solute pumping (Tab. 1). The baseline and pump-induced controls set *J*_poly_ = *γ* = 0; the actin-driven and combined conditions use positive *J*_poly_ and *γ* = 0.01 s^*−*1^. Active pumps are off in the baseline and actin-driven conditions and on in the pump-induced and combined conditions. The initial actin state and passive mechanical and transport coefficients are identical across the four conditions. F-actin network stress, all drag terms, membrane elasticity, and passive transport remain active (Table 1). The stress and drag forces evolve with the computed F-actin concentration, velocities, and membrane geometry. Their laboratory-frame displacements are denoted *d*_0_, *d*_*A*_, *d*_*W*_, and *d*_*AW*_. If the two active mechanisms were dynamically independent, the combined displacement would equal the baseline-corrected sum *d*_add_ = *d*_*A*_ + *d*_*W*_ *− d*_0_. We define the excess displacement as Δ*d* = *d*_*AW*_ *− d*_add_. The contrast has the symmetric form Δ*d* = (*d*_*AW*_ *− d*_*W*_) *−* (*d*_*A*_ *− d*_0_) = (*d*_*AW*_ *− d*_*A*_) *−* (*d*_*W*_ *− d*_0_). The first form compares the total response to actin turnover with pumps active and inactive; the second compares the total response to active solute pumping with membrane polymerization and bulk depolymerization both active or both inactive. Positive Δ*d* quantifies nonlinear coupling between active solute pumping and actin dynamics, including their coupled effects on actin transport, F-actin network stress, drag, fluid flow, and cell geometry.

**Table 1:**
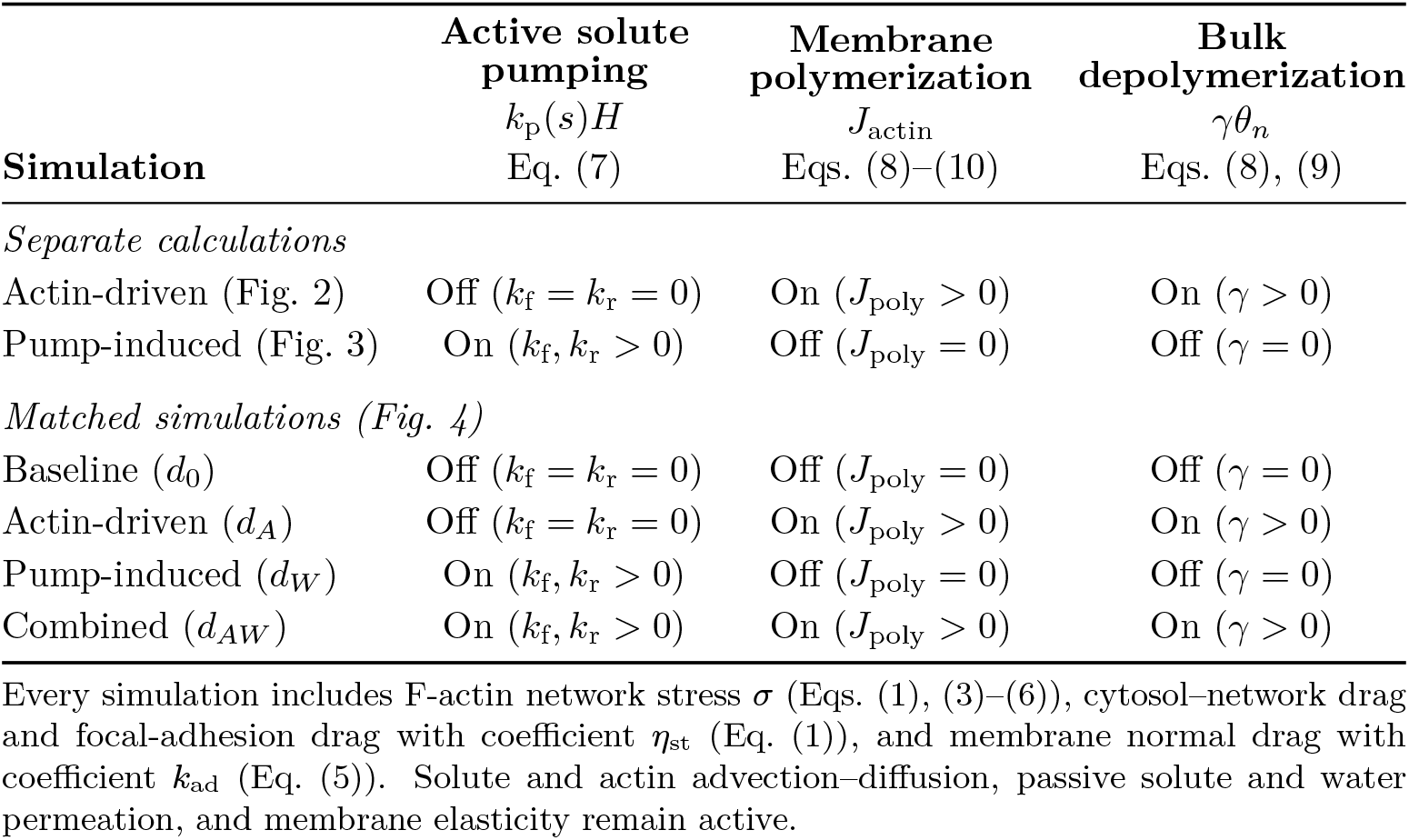
Processes varied across the simulations. On and Off denote enabled and suppressed terms.

| **Simulation** | **Active solute pumping**<br>*k*<sub>p</sub>(*s*)*H*<br>Eq. (7) | **Membrane polymerization**<br>*J*<sub>actin</sub><br>Eqs. (8)–(10) | **Bulk depolymerization**<br>*γθ*<sub>*n*</sub><br>Eqs. (8), (9) |
|---|---|---|---|
| *Separate calculations* |  |  |  |
| Actin-driven (Fig. 2) | Off (*k*<sub>f</sub> = *k*<sub>r</sub> = 0) | On (*J*<sub>poly</sub> > 0) | On (*γ* > 0) |
| Pump-induced (Fig. 3) | On (*k*<sub>f</sub>, *k*<sub>r</sub> > 0) | Off (*J*<sub>poly</sub> = 0) | Off (*γ* = 0) |
| *Matched simulations (Fig. 4)* |  |  |  |
| Baseline (*d*<sub>0</sub>) | Off (*k*<sub>f</sub> = *k*<sub>r</sub> = 0) | Off (*J*<sub>poly</sub> = 0) | Off (*γ* = 0) |
| Actin-driven (*d*<sub>*A*</sub>) | Off (*k*<sub>f</sub> = *k*<sub>r</sub> = 0) | On (*J*<sub>poly</sub> > 0) | On (*γ* > 0) |
| Pump-induced (*d*<sub>*W*</sub>) | On (*k*<sub>f</sub>, *k*<sub>r</sub> > 0) | Off (*J*<sub>poly</sub> = 0) | Off (*γ* = 0) |
| Combined (*d*<sub>*AW*</sub>) | On (*k*<sub>f</sub>, *k*<sub>r</sub> > 0) | On (*J*<sub>poly</sub> > 0) | On (*γ* > 0) |

Every simulation includes F-actin network stress *σ* (Eqs. (1), (3)–(6)), cytosol–network drag and focal-adhesion drag with coefficient *η*<sub>st</sub> (Eq. (1)), and membrane normal drag with coefficient *k*<sub>ad</sub> (Eq. (5)). Solute and actin advection–diffusion, passive solute and water permeation, and membrane elasticity remain active.

## 3 Results

### 3.1 Actin turnover and transport shape migration and cell morphology

Actin-driven migration is first examined with active solute pumping suppressed, *k*_f_ = *k*_r_ = 0. The three branches in the parameter sweeps correspond to three membrane normal-resistance coefficients, *k*_ad_. Over the sampled range, increasing the prescribed membrane polymerization capacity increases the computed migration velocity (Fig. 2A). Across the same sweep, smaller *k*_ad_ gives larger velocities, consistent with reduced normal resistance to membrane motion.

**Figure 2:**
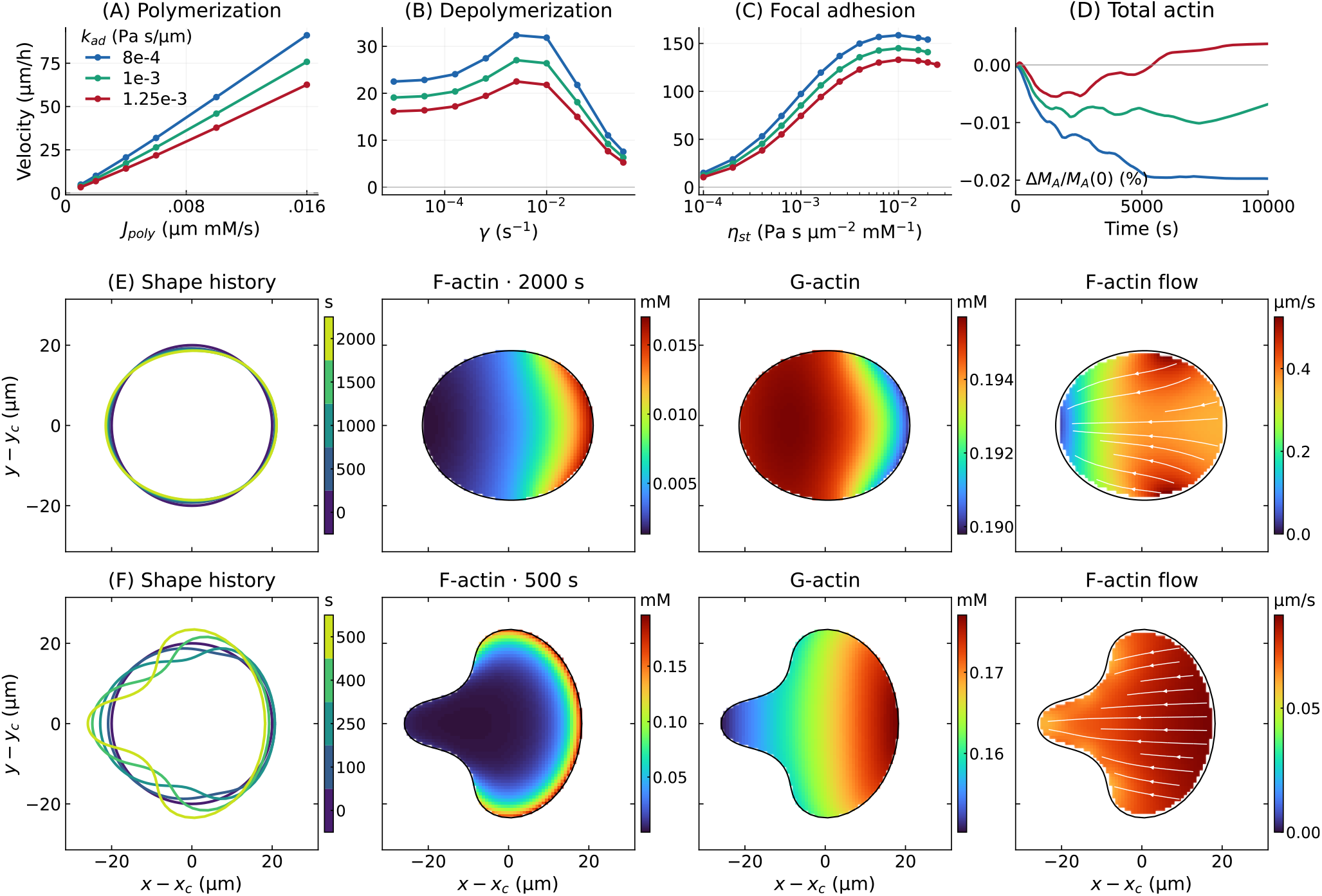
Actin-driven migration, morphology, and long-time conservation on a deforming domain. Active solute pumping is disabled. Blue, green, and red curves in A–D correspond to *k*_ad_ = 8 *×*10^*−*4^, 1 *×* 10^*−*3^, and 1.25 *×* 10^*−*3^ Pa*·*s/µm, respectively. Panels A–D use *θ*_*c,c*_ = 0.02 mM. **(A)** Migration velocity evaluated at 450 s from the centroid displacement while *J*_poly_ is varied at *γ* = 0.01 s^*−*1^ and *η*_st_ = 2 *×* 10^*−*4^ Pa*·* s/(µm^2^ *·*mM). **(B)** Migration velocity as *γ* is varied at *J*_poly_ = 6 *×* 10^*−*3^ µm*·* mM/s and the same *η*_st_. **(C)** Migration velocity as *η*_st_ is varied at *J*_poly_ = 6 *×* 10^*−*3^ µm*·*mM/s and *γ* = 10^*−*3^ s^*−*1^. The curves exhibit a shallow maximum near *η*_st_ = 10^*−*2^ Pa*·*s/(µm^2^ *·*mM) over the sampled range. **(D)** Relative change in total intracellular actin, *M*_*A*_ = *M*_*F*_ +*M*_*G*_, through 10,000 s for *J*_poly_ = 4 *×* 10^*−*3^ µm*·*mM/s, *γ* = 0.01 s^*−*1^, and *η*_st_ = 2 *×* 10^*−*4^ Pa s/(µm^2^ *·*mM). The absolute relative conservation error remains below 0.02% for all three *k*_ad_ values. **(E)** Shape history and F-actin, G-actin, and cell-relative F-actin velocity at 2000 s for *γ* = 0.04 s^*−*1^ and *η*_st_ = 2 *×* 10^*−*4^ Pa*·*s/(µm^2^ *·*mM). **(F)** Corresponding quantities at 500 s for *γ* = 0.01 s^*−*1^ and *η*_st_ = 2 *×*10^*−*2^ Pa*·*s/(µm^2^ *·*mM). E and F use *k*_ad_ = 10^*−*3^ Pa*·*s/µm, *J*_poly_ = 6*×*10^*−*3^ µm*·*mM/s, and *θ*_*c,c*_ = 2*×*10^*−*4^ mM. Numerical settings are given in Supplementary Sections S1–S4.

Depolymerization produces a biphasic cell velocity (Fig. 2B). A reported lamellipodial turnover time of approximately 61 s corresponds to an effective *γ≈* 0.016 s^*−*1^ [41], within the 0.01–0.04 s^*−*1^ range used in the field examples. At small *γ*, F-actin persists over a large fraction of the cell and the retrograde network flow is weak. Restoration of peripheral assembly by G-actin injection after cofilin inactivation [42] identifies monomer supply as the link between bulk disassembly and membrane assembly, represented by the *γθ*_*n*_ source in Eq. (9). Increasing *γ* strengthens turnover and retrograde flow, which raises migration speed. At larger *γ*, the effective spatial extent of the F-actin network decreases sufficiently that the reduction in the force-generating region outweighs the gain in retrograde flow, and the cell slows [26, 27]. The F-actin mass satisfies d*M*_*F*_ */*d*t* = ∫_Ω_ *J*_actin_ d*ℓ − γM*_*F*_. At steady inventory, *γM*_*F*_ ≤ ∫_Ω_ *j*(*s*) d*ℓ*; a larger depolymerization rate requires a larger membrane assembly flux to sustain a given F-actin mass. The maximum reflects the competition between filament lifetime, network motion, and the spatial extent over which substrate forces are generated.

Substrate coupling is also biphasic (Fig. 2C). At low *η*_st_, increasing substrate drag strengthens transmission of actin-generated force to the substrate and raises migration speed. Keratocytes on weakly adhesive substrates exhibit faster retrograde actin flow and slower cell translation [43], the paired response expected when network slip limits force transmission on this branch. The same parameter also reduces the fraction *a* = *η/*(*η* + *η*_st_) of cytosolic motion inherited by F-actin and lowers its effective diffusivity *D*_*n*_ = *k*_*σn*_*/*(*η* + *η*_st_). At larger *η*_st_, network motion is increasingly suppressed and F-actin redistribution becomes more localized, so the gain from stronger substrate coupling is offset by changes in retrograde flow and the spatial distribution of traction. This two-dimensional response is consistent with the biphasic focal-adhesion dependence obtained in our earlier one-dimensional G-actin model, where strong adhesion reduced the propulsive contribution of actin flow and altered the F-actin distribution [25]. The sampled range already resolves the maximum and extends into the regime in which *D*_*n*_ is small and network redistribution is strongly restricted. Migration responses to changes in ligand density, integrin abundance, and affinity collapsed onto a common dependence on measured cell–substratum adhesion strength [44]. In Eq. (1), this coupling is *η*_st_*θ*_*n*_***v***_*n*_, so receptor-level adhesion changes are represented by an effective drag coefficient per unit F-actin concentration.

The total actin conservation calculation tests the numerical treatment of boundary motion, membrane polymerization, bulk depolymerization, and transport of the two actin fields on the same deforming domain (Fig. 2D). The relative error remains below 0.02% through 10,000 s, which verifies the integrated material balance in these tests.

The actin depolymerization rate and substrate focal adhesion strength affect cell morphology and the spatial field of actin dynamics. For *γ* = 0.04 s^*−*1^, the characteristic filament lifetime is about 25 s, while *η*_st_ = 2 *×* 10^*−*4^ Pa*·*s/(µm^2^*·*mM) gives *D*_*n*_*≈* 5 µm^2^/s. The corresponding diffusive–turnover length 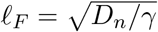 is about 11 µm before advection is included. F-actin assembled at the front can redistribute over a substantial fraction of the cell before depolymerizing. A smooth front-to-rear F-actin gradient develops, the complementary G-actin pool remains broadly distributed, and the F-actin flow is organized mainly along the migration axis and decreases towards the cell rear (Fig. 2E).

Reducing the depolymerization rate to 0.01 s^*−*1^ extends the filament lifetime to about 100 s, while increasing *η*_st_ by two orders of magnitude lowers *D*_*n*_ to approximately 0.05 µm^2^/s. The diffusive–turnover length consequently falls to about 2.2 µm. Newly assembled F-actin remains concentrated near the regions where it is produced, and the network velocity remains relatively uniform across the cell (Fig. 2F). Because the F-actin network stress is proportional to local F-actin concentration, this heterogeneous actin field produces heterogeneous membrane loading. The resulting lobes and constrictions arise from the coupled evolution of actin transport, network motion, cytosolic mechanics, and membrane shape. These actin-driven calculations provide the mechanical reference for the combined simulations.

### 3.2 Polarized active solute pumps generate osmotic flow and migration

Active solute transport causes asymmetric membrane flux, resulting in asymmetric intracellular solute concentration and osmotic pressure difference, *RT* (*c*_*i*_ *− c*_*e*_), along the cell membrane. The resulting osmotic pressure asymmetry drives normal water flux and intracellular flow. Increasing either pump strength or the spatial extent of the active profile increases the prescribed membrane transport and produces larger cell displacement (Fig. 3A).

**Figure 3:**
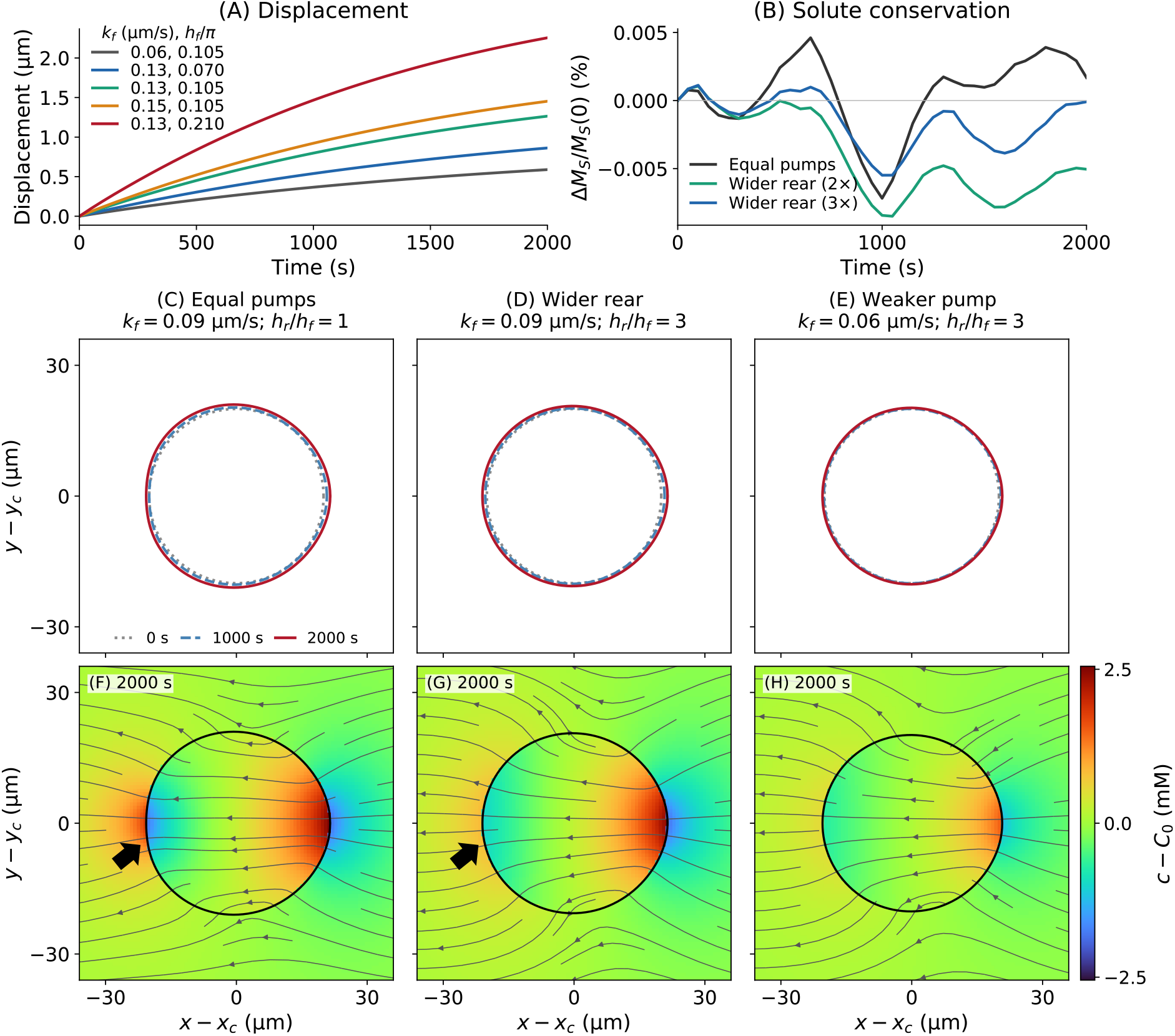
Pump-induced osmotic migration and solute conservation on a deforming permeable boundary. Pump-induced calculations begin from *F*_0_ = *G*_0_ = 0.1 mM and use *J*_poly_ = 0, *γ* = 0. **(A)** Laboratory-frame centroid displacement for five equal front/rear pump profiles; the front coefficient *k*_f_ and width *h*_f_*/π* are indicated in the legend, with *k*_r_ = *k*_f_ and *h*_r_ = *h*_f_. **(B)** Relative change in total solute at fixed front profile *k*_f_ = 0.09 µm/s and *h*_f_ = 0.105*π* for equal pumps, a twofold-wider rear profile, and a threefold-wider rear profile with the rear coefficient reduced so that the nominal integrated rear pumping capacity remains comparable. The absolute relative conservation error remains below 0.01% over 2000 s. **(C**,**F)** Equal pumps: *k*_f_ = *k*_r_ = 0.09 µm/s and *h*_f_ = *h*_r_ = 0.105*π*. **(D**,**G)** Wider rear: *k*_f_ = 0.09, *k*_r_ = 0.03 µm/s, *h*_f_ = 0.105*π*, and *h*_r_ = 0.315*π*. **(E**,**H)** Weaker broad-rear profile: *k*_f_ = 0.06, *k*_r_ = 0.02 µm/s with the same widths as D,G. C–E show cell shapes at 0, 1000, and 2000 s; F–H show *c − C*_0_ and cell-relative cytosolic-velocity streamlines at 2000 s. Numerical settings are given in Supplementary Sections S1–S4.

The solute calculation provides an independent test of the sharp moving-boundary transport solver (Fig. 3B). Solute crosses the membrane through matched one-sided Robin conditions while osmotic water flux continuously changes the geometry. The absolute relative solute error remains below 0.01% for all three pump profiles in this test. Together with the actin test, this result demonstrates high-precision conservation for two distinct transport problems: transfer across a permeable moving interface and interconversion of two intracellular fields transported at different velocities.

The equal-pump and broad-rear cases separate the amount of prescribed rear activity from its localization (Fig. 3C,D,F,G). In the equal-pump case, *k*_f_ = *k*_r_ and *h*_f_ = *h*_r_. In the broad-rear case, the rear coefficient is reduced threefold while its width is increased threefold, so *k*_r_*h*_r_ remains comparable to *k*_f_*h*_f_. Our model illustrates that solute flux is not determined by this product alone because the boundary condition contains *k*_p_*H* and the donor concentration *H* evolves with the cell. Spreading the rear pump changes where solute is removed, which reshapes the concentration field and the local osmotic pressure that drives water through the membrane (Fig. 3F,G). Optogenetic redistribution of SWELL1 halted confined migration when its abundance was equalized between the front and rear of the cell and reversed migration when SWELL1 became enriched at the front [22]. In the model, the spatial distribution of *k*_p_*H* sets osmotic loading and water flux. The physical length profile is further influenced by membrane stretch d*ℓ* = *J* d*s*.

The resulting streamline is consistent with the solute fields: solute accumulates toward the front and is depleted toward the rear, producing water entry at the front, water exit at the rear, and a predominantly rearward cytosolic flow in the cell frame (Fig. 3F–H). The broader rear profile distributes solute export over a larger portion of the posterior membrane at comparable nominal integrated rear transport capacity (Fig. 3C,D,F,G). Reducing both active solute pump coefficients while retaining the broad-rear profile weakens the concentration contrast, cell-relative flow, and displacement (Fig. 3E,H).

The model links *k*_f_ to cell motion through intracellular osmotic pressure and water uptake. The reduced swelling and migration observed after NHE1 depletion in confined MDA-MB-231 cells [22] are consistent with the coupled effects expected from reduced solute uptake. Similarly, the suppression of confined migration following AQP5 depletion [20] independently links motility to the water conductance *k*_w_, which translates osmotic drive into membrane motion.

### 3.3 Coupled hydraulic and actin mechanics enhance migration

The actin kinetic parameters and passive material coefficients remain fixed across the pump-strength and pump-width series. Cell geometry, water permeation, actin concentrations, and network motion evolve together through the shared fluid–membrane and transport equations.

Increasing pump strength at fixed width and increasing pump width at fixed peak coefficient change both cell size and the spatial actin distributions (Fig. 4A,B). F-actin remains enriched toward the front, whereas G-actin develops a rear bias (Fig. 4C,D). The increasing normalized F-actin polarity describes a stronger relative front–rear concentration contrast within the changing cell geometry.

**Figure 4:**
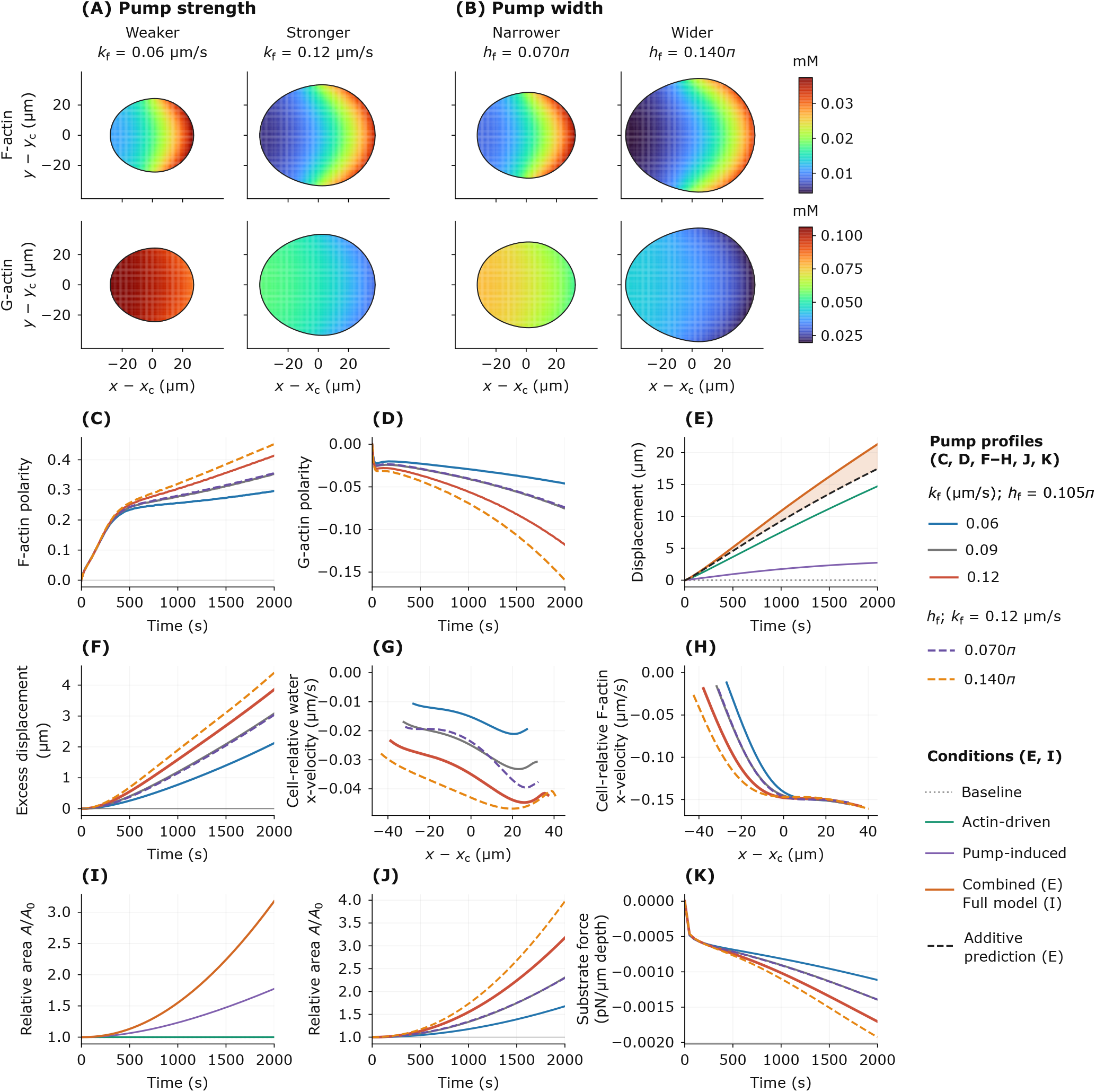
Coupled actin redistribution, migration, and force transmission. **(A**,**B)** F-actin (upper) and G-actin (lower) concentrations at 2000 s for changes in pump strength and width, respectively. Coordinates are relative to the area centroid; the front is at positive *x*. **(C**,**D)** F-actin and G-actin polarity; positive values denote front enrichment and negative values rear enrichment. **(E)** Laboratory-frame centroid displacements for baseline (*d*_0_), actin-driven (*d*_*A*_), pump-induced (*d*_*W*_), and combined (*d*_*AW*_; labeled Full model in I) conditions, together with the baseline-corrected sum *d*_add_ = *d*_*A*_ + *d*_*W*_ *− d*_0_. **(F)** Excess displacement Δ*d* = *d*_*AW*_ *− d*_add_ for the five pump profiles. **(G**,**H)** Cytosolic and F-actin *x*-velocities relative to the cell along the horizontal centroid line at 2000 s. **(I**,**J)** Relative enclosed area *A*(*t*)*/A*_0_, where *A*_0_ = *A*(0), for the four conditions and five combined pump profiles, respectively. **(K)** Horizontal force exerted on the substrate by distributed actin–substrate drag, per unit out-of-plane thickness; negative values denote rearward force. Membrane normal resistance contributes separately. All pump profiles use *k*_r_ = *k*_f_*/*3 and *h*_r_ = 3*h*_f_. Membrane polymerization and bulk depolymerization are inactive in the baseline and pump-induced controls (*J*_poly_ = *γ* = 0) and active in the actin-driven and combined conditions. Pumps are active in the pump-induced and combined conditions. F-actin network stress and drag remain active in all conditions. Parameters and diagnostic definitions are given in Supplementary Sections S1, S4, and S5.

The matched controls quantify the net interaction between actin dynamics and active solute pumps. After an initial transient, the combined displacement exceeds the baseline-corrected sum of the isolated active responses (Fig. 4E). The excess displacement Δ*d* remains positive as active solute pumps become stronger or more spatially extended (Fig. 4F). Grid, membrane, and time-step refinement retain non-trivial excess displacement. The common comparison intervals and the late-time sensitivity of displacement, area, and substrate force are reported in Supplementary Section S4 (Supplementary Fig. S1).

Stronger active solute pumps produce a more rearward cell-relative cytosolic velocity, while the F-actin profiles reach comparable rearward speeds across the pump series (Fig. 4G,H). The cell-relative network velocity is 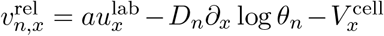, where 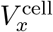 is the laboratory-frame centroid velocity (Eq. (2)). As the cell expands, retrograde F-actin motion extends over a longer physical span along the migration axis.

The area histories show that the baseline and actin-driven cells remain close to their initial area, whereas the pump-induced and combined cells expand, with greater expansion in the combined condition (Fig. 4I). Incompressibility and the membrane kinematics give d*A/*d*t* = ∫_Ω_ *w* d*ℓ*, so area growth measures net water entry. Actin conservation gives a mean total concentration *M*_*A*_*/A* that decreases as the cell expands, while polymerization and depolymerization change the partition between the F-actin and G-actin pools. At 2000 s, the widest pump profile gives both the largest area increase and the largest excess displacement (Fig. 4F,J). Increasing the strength or spatial extent of active solute pumps also produces a more rearward horizontal force on the substrate (Fig. 4K). We calculate this force per unit out-of-plane thickness as 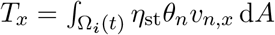. Negative *T*_*x*_ denotes rearward traction on the substrate and a forward reaction on the actin network. We reconstruct the integral from the laboratory-frame network flux, as detailed in Supplementary Section S5.

## 4 Discussion and Conclusion

Our earlier energetic analyses treated actin polymerization and active solute pumping as distinct sources of free-energy input [24]. In the present work, we resolve their collective mechanical output as cell geometry, water permeation, and actin dynamics adjust together. The nonlinear excess displacement reflects coupled changes in cell morphology, net water retention, actin motion, and substrate force transmission. Actin polymerization deforms the cell, and its interaction with active solute pumps produces further changes in morphology and cell area. Area growth records accumulated net water entry at fixed hydraulic permeability. The expanding cell supports retrograde F-actin motion across a longer physical span, and the integrated rearward force on the substrate increases in magnitude. The corresponding forward reaction on the actin network contributes to the collective mechanical response that supports enhanced migration (Fig. 5).

**Figure 5:**
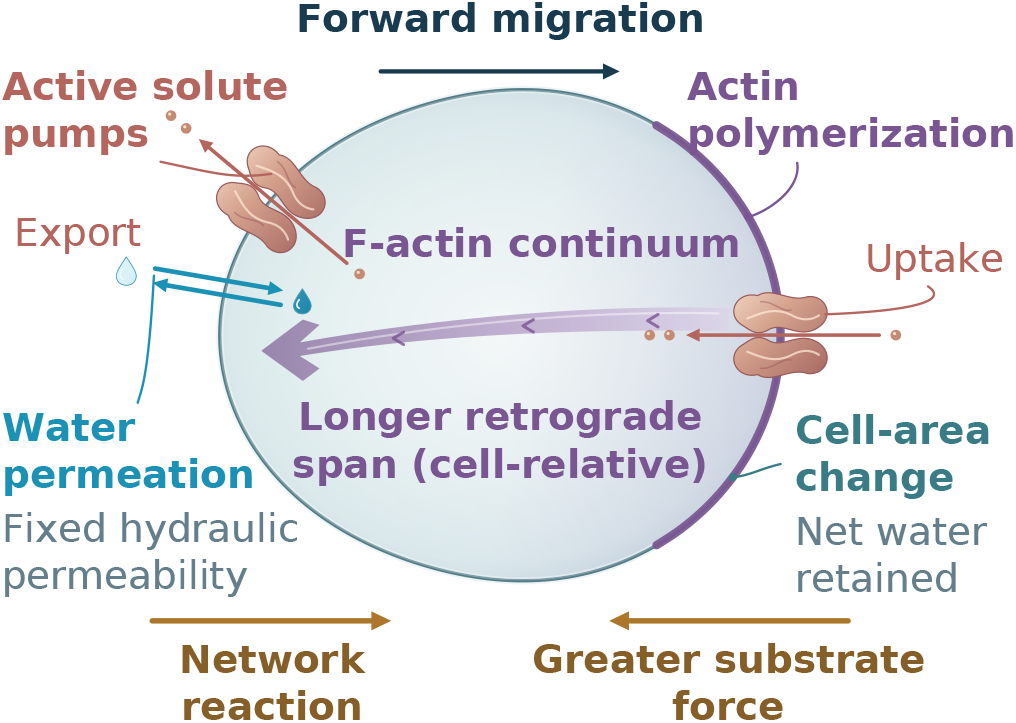
Coupled actin mechanics and active solute transport. A schematic plan view illustrates the membrane-bounded cell and its mechanical coupling to the stationary substrate. Leading-edge actin polymerization generates membrane deformation through F-actin network stress, while active solute pumps transfer neutral solute across the same deforming membrane. Purple shading denotes the continuum F-actin phase, the boundary band denotes actin assembly, and the broad purple arrow indicates cell-relative retrograde network motion over an extended cell span. Generic coral membrane symbols denote active solute pumps and the associated solute transfer. The bidirectional blue symbol denotes water permeation at fixed hydraulic permeability; cell-area change records accumulated net water retention. Gold arrows show equal and opposite whole-cell drag resultants: rearward force on the substrate and forward reaction on the network.

Resolving these coupled processes requires accurate mechanics and transport on a moving permeable boundary. Osmotic forcing depends on one-sided solute concentrations, while actin polymerization transfers material between two fields that move with different velocities. The present framework imposes the Robin conditions sharply on the accepted membrane geometry and applies the F-actin/G-actin conversion as paired interface fluxes. Independent long-time calculations show excellent conservation of total solute and total actin under sustained deformation and transport (Fig. 3B; Fig. 2D). In addition, we performed mesh-refinement comparisons to evaluate the sensitivity of displacement, area, and substrate force to numerical resolution (Supplementary Section S4).

The actin-driven calculations define how migration varies with actin turnover and substrate coupling and provide reference responses for the coupled system (Fig. 2B,C). In the pump-induced comparison, a broader rear profile with a proportionately lower peak changes the solute and flow distributions at comparable nominal rear capacity *k*_r_*h*_r_ (Fig. 3C,D,F,G). The local active flux depends on donor concentration through *k*_p_*H* (Eq. (7)). These results are consistent with experiments in which polarized ion transporters and aquaporins regulate the location of water entry and exit during migration [20, 22].

Osmotic permeabilities of approximately 100 and 290 µm/s were measured in AQP5- and AQP4-expressing Xenopus oocytes, respectively [45]. At the model value *RT* = 2.494*×*10^3^ Pa *·* m^3^/mol, 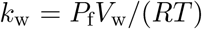 with water molar volume 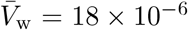 m^3^/mol gives approximately 0.72 *×* 10^*−*6^ and 2.1 *×* 10^*−*6^ µm/(Pa*·*s), comparable in scale to the 1 *×*10^*−*6^ and 3 *×*10^*−*6^ used here.

NHE1 overexpression partially restored motility in latrunculin-A-treated MDA-MB-231 cells at elevated extracellular viscosity [46]. In the model, the osmotic pressure jump *RT* [*c*] and F-actin network stress *σ* enter the same water-permeation law (Eq. (6)), providing a mechanical route for increased water uptake to sustain migration when actin assembly is reduced.

Setting *k*_f_ = *k*_r_ = 0 removes prescribed active solute transport while retaining passive solute and water permeation. Setting *J*_poly_ = 0 removes all membrane assembly; inhibition of one assembly pathway is represented by a partial capacity reduction with residual assembly, consistent with distinct formin- and Arp2/3-dependent monomer use [16]. Residual actin disassembly after myosin-II inhibition [47] supports a myosin-independent component of the depolymerization rate. Persistent migration after latrunculin treatment in 3 µm-wide channels [20] provides the biological setting in which membrane transport sustains motion after actin assembly is blocked.

The present formulation uses a neutral solute and a two-field representation of actin. Ion-specific electrodiffusion, explicit myosin dynamics, regulatory signaling, membrane bending, and three-dimensional geometry can be incorporated when required by the biological setting. Within this framework, enhanced migration emerges from the coupled evolution of actin dynamics, active solute transport, cell geometry, and substrate force transmission. Future experiments that simultaneously measure cell area and shape, actin motion, and substrate traction during perturbations of transporter activity or localization could test how hydraulic and mechanical responses coordinate to drive cell migration.

## Data and code availability

The computational package is publicly available through the [GitHub repository](https://github.com/han-zhou-math/simcell.git).

## Acknowledgments

Yizeng Li is supported by National Science Foundation (NSF) 2303648. The opinions, findings, and conclusions, or recommendations expressed are those of the authors and do not necessarily reflect the views of any of the funding agencies.

