## Supplementary Materials for "A conservative moving-boundary multiphysics framework reveals nonlinear coupling between osmotic flow and actin-driven cell migration"


### S1. Physical parameters and constitutive specifications

All dimensional parameter values used in the simulations are collected here. Pump strength denotes the coefficient in the membrane solute-transfer law and is distinct from the realized transmembrane solute flux. Pump widths are defined in the membrane material coordinate. The fluid viscosity is identical inside and outside the cell. The initial cell is circular with radius $2.00 \times 10^1$ μm, and the initial solute concentration is $3.40 \times 10^2$ mM on both sides of the membrane. The supplementary notation is $F = \theta_n$, $G = \theta_c$, $\boldsymbol{v}_F = \boldsymbol{v}_n$, $D_F = D_n$, $D_G = D_c$, $D_S = D_\mathrm{s}$, $k_\sigma = k_{\sigma n}$, and $k_c = k_\mathrm{sol}$. All reported migration calculations begin from $F = G = 1.00 \times 10^{-1}$ mM. Standalone pump-induced calculations set $J_\mathrm{poly} = \gamma = 0$, with membrane polymerization and bulk depolymerization inactive (Fig. 3). The matched baseline and pump-induced controls use $J_\mathrm{poly} = \gamma = 0$; the actin-driven and combined conditions use $J_\mathrm{poly} > 0$ and $\gamma > 0$ (Fig. 4; Table S3). Every calculation includes F-actin network stress $\sigma$ and drag terms with coefficients $\eta$, $\eta_\mathrm{st}$, and $k_\mathrm{ad}$.

F-actin transport follows

$$a = \frac{\eta}{\eta + \eta_\mathrm{st}}, \qquad D_F = \frac{k_\sigma}{\eta + \eta_\mathrm{st}}, \qquad \boldsymbol{v}_F = a\boldsymbol{u} - D_F \nabla \log F.$$

At $\eta_\mathrm{st} = 2.00 \times 10^{-4}\,\mathrm{Pa\,s/(\mu m^2\,mM)}$, the derived diffusivity is $D_F = 5.00\,\mathrm{\mu m^2/s}$. The membrane uses the linear membrane tension law $T_\mathrm{m} = k_\mathrm{m} J$ with modulus $k_\mathrm{m} = 1.00 \times 10^{-5}$ Pa. No independent bending force or explicit area penalty is active in the reported simulations. The membrane normal-resistance coefficient $k_\mathrm{ad}$ and the F-actin substrate-drag coefficient $\eta_\mathrm{st}$ are distinct parameters. Passive osmotic feedback remains present when active pumping is disabled.

Localized polymerization has capacity $J_\mathrm{poly} g(s)$ and saturation factor $G/(G + \theta_{c,c})$, with $g(s) = 2 - \tanh(s^6/d_\mathrm{w}) - \tanh[(2\pi - s)^6/d_\mathrm{w}]$ and $d_\mathrm{w} = 1.00$. The material coordinate $s$ labels membrane points and is dimensionless; $d_\mathrm{w}$ is not a physical width. Front and rear pump profiles are centered at $s = 0$ and $s = \pi$, respectively, with the initial leading direction along positive $x$. Pumps are active from the initial time in pump-enabled cases.


*

**Supplementary Table S1. Common physical parameters.**

| Symbol | Meaning | Value and unit | Source / justification |
|---|---|---|---|
| $\mu$ | Fluid dynamic viscosity | $1.00 \times 10^{-3}$ Pa s | Water-like cytosolic viscosity; the same scale is used in earlier migration models [1]. |
| $b_\mathrm{i}$ | Additional intracellular Brinkman drag | 0 Pa s/$\mu$m$^2$ | Model choice; no additional bulk porous drag is imposed. |
| $b_\mathrm{e}$ | Additional extracellular Brinkman drag | 0 Pa s/$\mu$m$^2$ | Model choice; no additional bulk porous drag is imposed. |
| $k_\mathrm{m}$ | Membrane elastic modulus | $1.00 \times 10^{-5}$ Pa | Free effective parameter of the two-dimensional linear membrane tension law; the constitutive form follows earlier two-dimensional models [2]. |
| $\ell_\mathrm{ref}$ | Reference material metric | 0 $\mu$m/rad | Zero reference stretch in the linear membrane tension law. |
| $\eta$ | Cytosol–network drag per actin concentration | $1.00 \times 10^{-8}$ Pa s/($\mu$m$^2$ mM) | Effective two-phase drag coefficient inherited from the interpenetrating-network formulation [2, 3]. |
| $k_\sigma$ | Coefficient of F-actin pressure | $1.00 \times 10^{-3}$ Pa/mM | Free effective coefficient in the present two-dimensional actin-stress law; the same constitutive structure is used previously [2]. |
| $D_G$ | G-actin diffusivity | $1.00 \times 10^{1}$ $\mu$m$^2$/s | Cytoplasmic G-actin diffusivities are of order 10 $\mu$m$^2$/s; direct measurements report 13.7 $\mu$m$^2$/s [4]. |
| $d_\mathrm{w}$ | Localization denominator in polymerization profile | 1.00 rad$^6$ | Free dimensionless profile-shape parameter in membrane material coordinates. |
| $F_\mathrm{load}$ | External force per out-of-plane depth | 0 Pa $\mu$m | Model choice; no external mechanical load is applied. |
| $D_S$ | Solute diffusivity | $1.00 \times 10^{2}$ $\mu$m$^2$/s | Small solutes diffuse in mammalian cytoplasm on the $10^2$ $\mu$m$^2$/s scale [5, 6]. |
| $k_\mathrm{c}$ | Passive membrane solute permeability | 1.00 $\mu$m/s | Free effective permeability for the neutral model solute; the same velocity-type coefficient is used in earlier osmotic-migration models [1]. |
| $C_0$ | Initial solute concentration inside and outside | $3.40 \times 10^{2}$ mM | Physiological osmolarity scale used in osmotic-engine experiments and models [7]. |
| $\nabla C_0$ | Initial opposing solute gradient | 0 mM/$\mu$m | Initial condition; osmotic gradients develop dynamically from membrane transport. |
| $R_0$ | Initial cell radius | $2.00 \times 10^{1}$ $\mu$m | Typical mammalian-cell scale and consistent with earlier two-dimensional calculations [1, 2]. |
| $q_0$ | Initial axis ratio | 1.00 | Circular initial cell. |
| $s_0$ | Initial shape perturbation | 0 | No imposed shape perturbation. |
| $x_0, y_0$ | Initial center offsets | 0 $\mu$m | Coordinate choice. |
| $RT$ | Effective osmotic coefficient | $2.494 \times 10^{3}$ Pa/mM | $RT$ at 300 K. |
| $k_\mathrm{b}$ | Bending coefficient | 0 | No independent bending contribution is active. |
| $F_0$ | Initial F-actin concentration | $1.00 \times 10^{-1}$ mM | Together with $G_0$, gives a total actin scale consistent with earlier actin models and experimental concentration estimates [8, 9]. |

| Symbol | Meaning | Value and unit | Source / justification |
|---|---|---|---|
| $G_0$ | Initial G-actin concentration | $1.00 \times 10^{-1}$ mM | Same justification as $F_0$; this common partition is used in actin-driven, pump-induced, baseline, and coupled calculations. |

### S1.1. Parameters varied between simulations

#### Supplementary Table S2. Parameters varied between simulations.

| Symbol | Meaning | Values used | Source / justification |
|---|---|---|---|
| $k_{\mathrm{ad}}$ | Membrane normal resistance | Fig. 2 A–D: $8.00 \times 10^{-4}$, $1.00 \times 10^{-3}$, $1.25 \times 10^{-3}$ Pa s/µm; Fig. 2 E,F and Figs. 3–4: $1.00 \times 10^{-3}$ Pa s/µm | Effective drag coefficient for normal membrane motion relative to the stationary substrate; the permeable moving-boundary form follows earlier osmotic formulations [10]. |
| $\eta_{\mathrm{st}}$ | Substrate drag per actin concentration | Fig. 2C: $1.00 \times 10^{-4}$–$2.50 \times 10^{-2}$ Pa s/(µm$^2$ mM), with the plotted values listed in Table S3; Fig. 2A,B,D,E: $2.00 \times 10^{-4}$; Fig. 2F: $2.00 \times 10^{-2}$; Figs. 3–4: $2.00 \times 10^{-4}$ | Free effective drag parameter. Traction stress varies biphasically with actin retrograde flow speed [11]; the effective substrate-drag law follows earlier migration models [1]. |
| $\gamma$ | Depolymerization rate | Fig. 2B: $1.00 \times 10^{-5}$–$3.20 \times 10^{-1}$ s$^{-1}$; Fig. 2C: $1.00 \times 10^{-3}$ s$^{-1}$; Fig. 2A,D,F: $1.00 \times 10^{-2}$ s$^{-1}$; Fig. 2E: $4.00 \times 10^{-2}$ s$^{-1}$; Fig. 3: 0; Fig. 4: 0 in the baseline and pump-induced controls; $1.00 \times 10^{-2}$ s$^{-1}$ in the actin-driven and combined conditions | Turnover range follows the earlier actin-depolymerization studies [2, 8]. At $\gamma = 0$, both the F-actin sink and G-actin source vanish. |
| $J_{\mathrm{poly}}$ | Polymerization capacity | Fig. 2A: $1.00 \times 10^{-3}$–$1.60 \times 10^{-2}$ µm mM/s; Fig. 2B,C,E,F: $6.00 \times 10^{-3}$; Fig. 2D: $4.00 \times 10^{-3}$; Fig. 3: 0; Fig. 4: 0 in baseline/pump-induced controls and $6.00 \times 10^{-3}$ in actin/combined controls | The $6.00 \times 10^{-3}$ µm mM/s scale follows the earlier G-actin model; broader values provide sensitivity tests [8]. |
| $\theta_{c,c}$ | G-actin saturation concentration | Fig. 2A–D: $2.00 \times 10^{-2}$ mM; Fig. 2E,F and the actin-driven and combined conditions in Fig. 4: $2.00 \times 10^{-4}$ mM | The smaller value follows the actin critical-concentration scale used previously; the larger value reduces saturation and is used as a sensitivity setting [8, 9]. |
| $k_{\mathrm{w}}$ | Hydraulic permeability | Figs. 2–3: $1.00 \times 10^{-6}$ µm/(Pa s); Fig. 4: $3.00 \times 10^{-6}$ µm/(Pa s) | These values correspond to osmotic permeabilities within the range measured for aquaporin-expressing membranes [12, 13]. |
| $k_{\mathrm{f}}$ | Front active-pump coefficient | Fig. 3: $6.00 \times 10^{-2}$–$1.50 \times 10^{-1}$ µm/s; Fig. 4: 0, $6.00 \times 10^{-2}$, $9.00 \times 10^{-2}$, $1.20 \times 10^{-1}$ µm/s | Free effective active-transport coefficient; polarized ion and water transport is supported by osmotic-engine experiments [7]. |
| $h_{\mathrm{f}}$ | Front pump width | Fig. 3: $0.070\pi$, $0.105\pi$, $0.210\pi$; Fig. 4: $0.070\pi$, $0.105\pi$, $0.140\pi$ | Free spatial-profile parameter in membrane material coordinates. |
| $k_{\mathrm{r}}$ | Rear active-pump coefficient | Fig. 3: values paired with the front profiles, including $9.00 \times 10^{-2}$, $4.50 \times 10^{-2}$, $3.00 \times 10^{-2}$, and $2.00 \times 10^{-2}$ µm/s; Fig. 4: $k_{\mathrm{r}} = k_{\mathrm{f}}/3$ in pump-enabled cases | Free effective active-transport coefficient; rear efflux polarity is consistent with confined-migration experiments [7]. |

| | | | |
|---|---|---|---|
| $h_r$ | Rear pump width | Fig. 3: $h_r/h_f = 1$, 2, or 3; Fig. 4: $h_r = 3h_f$ in pump-enabled cases | Free spatial-profile parameter used to separate transport strength from localization. |

#### S1.2. Panel-specific parameter combinations

**Supplementary Table S3. Main parameter combinations used in the figures.**

| Panel | Varied quantity | Fixed active parameters and notes |
|---|---|---|
| 2A | $J_{\mathrm{poly}} = 1.00 \times 10^{-3}$, $2.00 \times 10^{-3}$, $4.00 \times 10^{-3}$, $6.00 \times 10^{-3}$, $1.00 \times 10^{-2}$, $1.60 \times 10^{-2}$ µm mM/s | $\gamma = 1.00 \times 10^{-2}$ s$^{-1}$; $\eta_{\mathrm{st}} = 2.00 \times 10^{-4}$ Pa s/(µm$^2$ mM); $\theta_{c,c} = 2.00 \times 10^{-2}$ mM; $k_{\mathrm{ad}} = 8.00 \times 10^{-4}$, $1.00 \times 10^{-3}$, $1.25 \times 10^{-3}$ Pa s/µm. |
| 2B | $\gamma = 1.00 \times 10^{-5}$, $4.00 \times 10^{-5}$, $1.60 \times 10^{-4}$, $6.40 \times 10^{-4}$, $2.56 \times 10^{-3}$, $1.00 \times 10^{-2}$, $4.00 \times 10^{-2}$, $1.60 \times 10^{-1}$, $3.20 \times 10^{-1}$ s$^{-1}$ | $J_{\mathrm{poly}} = 6.00 \times 10^{-3}$ µm mM/s; $\eta_{\mathrm{st}} = 2.00 \times 10^{-4}$; same $\theta_{c,c}$ and $k_{\mathrm{ad}}$ values as 2A. |
| 2C | $\eta_{\mathrm{st}} = 1.00 \times 10^{-4}$, $2.00 \times 10^{-4}$, $4.00 \times 10^{-4}$, $6.325 \times 10^{-4}$, $1.00 \times 10^{-3}$, $1.60 \times 10^{-3}$, $2.50 \times 10^{-3}$, $4.00 \times 10^{-3}$, $6.30 \times 10^{-3}$, $1.00 \times 10^{-2}$, $1.60 \times 10^{-2}$, $2.00 \times 10^{-2}$ Pa s/(µm$^2$ mM); the $k_{\mathrm{ad}} = 1.25 \times 10^{-3}$ branch also includes $2.50 \times 10^{-2}$ | $J_{\mathrm{poly}} = 6.00 \times 10^{-3}$ µm mM/s; $\gamma = 1.00 \times 10^{-3}$ s$^{-1}$; $\theta_{c,c} = 2.00 \times 10^{-2}$ mM. |
| 2D | Conservation through $1.00 \times 10^4$ s | $J_{\mathrm{poly}} = 4.00 \times 10^{-3}$ µm mM/s; $\gamma = 1.00 \times 10^{-2}$ s$^{-1}$; $\eta_{\mathrm{st}} = 2.00 \times 10^{-4}$; same three $k_{\mathrm{ad}}$ values. |
| 2E | Field example at $2.00 \times 10^3$ s | $J_{\mathrm{poly}} = 6.00 \times 10^{-3}$; $\gamma = 4.00 \times 10^{-2}$ s$^{-1}$; $\eta_{\mathrm{st}} = 2.00 \times 10^{-4}$; $\theta_{c,c} = 2.00 \times 10^{-4}$ mM; $k_{\mathrm{ad}} = 1.00 \times 10^{-3}$ Pa s/µm. |
| 2F | Field example at $5.00 \times 10^2$ s | $J_{\mathrm{poly}} = 6.00 \times 10^{-3}$; $\gamma = 1.00 \times 10^{-2}$ s$^{-1}$; $\eta_{\mathrm{st}} = 2.00 \times 10^{-2}$; $\theta_{c,c} = 2.00 \times 10^{-4}$ mM; $k_{\mathrm{ad}} = 1.00 \times 10^{-3}$ Pa s/µm. |
| 3A | Equal front/rear pump profiles | $(k_f, h_f/\pi) = (0.06, 0.105)$, $(0.13, 0.070)$, $(0.13, 0.105)$, $(0.15, 0.105)$, $(0.13, 0.210)$, with $k_r = k_f$ and $h_r = h_f$. |
| 3B | Rear-pump localization at fixed front profile | $k_f = 0.09$ µm/s, $h_f = 0.105\pi$; equal rear, twofold wider rear with half coefficient, and threefold wider rear with one-third coefficient. |
| 3C,F | Equal pumps | $k_f = k_r = 0.09$ µm/s; $h_f = h_r = 0.105\pi$. |
| 3D,G | Wider rear | $k_f = 0.09$, $k_r = 0.03$ µm/s; $h_f = 0.105\pi$, $h_r = 0.315\pi$. |
| 3E,H | Weaker broad rear | $k_f = 0.06$, $k_r = 0.02$ µm/s; same widths as 3D,G. |
| 4A,C,D,F–H,J,K | Pump-strength series | Profiles K06, K09, and K12 in Table S4; the field maps use K06 and K12. |
| 4B,C,D,F–H,J,K | Pump-width series | Profiles H070, K12, and H140 in Table S4; the field maps use H070 and H140. |
| 4E,I | Matched-control displacement and relative area | Pump-enabled conditions: $k_f = 0.12$, $k_r = 0.04$ µm/s; $h_f = 0.105\pi$, $h_r = 0.315\pi$. Pumps are off in the baseline and actin-driven conditions. All four conditions share the same initial $F_0 = G_0 = 0.1$ mM. The baseline and pump-induced controls use $J_{\mathrm{poly}} = \gamma = 0$. The actin-driven and combined conditions use $J_{\mathrm{poly}} = 6.00 \times 10^{-3}$ µm·mM/s and $\gamma = 1.00 \times 10^{-2}$ s$^{-1}$. |

### S2. Cartesian fluid grid and immersed membrane

The computational box is periodic in the migration direction and has stationary no-slip boundaries at the upper and lower walls. Pressure and concentrations are stored at cell centers on a uniform Cartesian mesh; the two fluid-velocity components are stored on the corresponding cell faces in a standard MAC arrangement. The membrane is discretized by Lagrangian markers $\boldsymbol{X}_k = \boldsymbol{X}(s_k)$, where $s_k = (k-1)\Delta s$

**Table S4:** Pump coefficients and material-coordinate widths for the coupled calculations.

| Profile | $k_\mathrm{f}$ (µm/s) | $k_\mathrm{r}$ (µm/s) | $h_\mathrm{f}/\pi$ | $h_\mathrm{r}/\pi$ |
|---|---|---|---|---|
| K06 | 0.06 | 0.02 | 0.105 | 0.315 |
| K09 | 0.09 | 0.03 | 0.105 | 0.315 |
| K12 | 0.12 | 0.04 | 0.105 | 0.315 |
| H070 | 0.12 | 0.04 | 0.070 | 0.210 |
| H140 | 0.12 | 0.04 | 0.140 | 0.420 |

and periodic indexing is used. Marker force spreading and velocity interpolation use the same four-point regularized delta kernel, so the interpolation operator is the discrete adjoint of force spreading [14].

For a grid function $w$, the finite-difference and averaging operators are

$$D_x^{\pm} w_{\alpha,\beta} = \pm\frac{w_{\alpha\pm1,\beta} - w_{\alpha,\beta}}{h}, \quad D_y^{\pm} w_{\alpha,\beta} = \pm\frac{w_{\alpha,\beta\pm1} - w_{\alpha,\beta}}{h}, \quad \Delta_h w = D_x^+ D_x^- w + D_y^+ D_y^- w, \tag{1}$$

$$A_x^+ w_{\alpha,\beta} = \tfrac{1}{2}(w_{\alpha+1,\beta} + w_{\alpha,\beta}), \qquad A_y^+ w_{\alpha,\beta} = \tfrac{1}{2}(w_{\alpha,\beta+1} + w_{\alpha,\beta}), \qquad D_t^- w^{n+1} = \frac{w^{n+1} - w^n}{\Delta t}. \tag{2}$$

The discrete membrane force is

$$\boldsymbol{F}_{\mathrm{mem},k} = k_\mathrm{m} D_s^+ D_s^- \boldsymbol{X}_k = \frac{k_\mathrm{m}}{\Delta s^2}\big(\boldsymbol{X}_{k+1} - 2\boldsymbol{X}_k + \boldsymbol{X}_{k-1}\big). \tag{3}$$

The fluid–membrane system uses a semi-implicit midpoint scheme. For a step from accepted state $n$ to updated state $n+1$, the midpoint marker positions and membrane displacement velocity are

$$\bar{\boldsymbol{X}}_k = \frac{\boldsymbol{X}_k^n + \boldsymbol{X}_k^{n+1}}{2}, \qquad \boldsymbol{V}_k^{n+1} = \frac{\boldsymbol{X}_k^{n+1} - \boldsymbol{X}_k^n}{\Delta t}.$$

The nonlinear membrane residual is treated by Newton iteration with matrix-free GMRES. The Jacobian approximation differentiates the elastic force while holding the interpolation kernel, normals, and metric fixed; the outer iteration recomputes the full nonlinear residual. Each Krylov product contains a Stokes–Brinkman solve; Fourier decomposition in the periodic direction reduces the fluid problem to banded transverse solves. The labels $\boldsymbol{u}^{n+1}$ and $p^{n+1}$ denote the fluid outputs of this midpoint-geometry solve. This construction follows the implicit immersed-boundary strategy used for elastic structures and the earlier osmotic moving-membrane formulation [10, 15].

The elastic force $\bar{\boldsymbol{F}}_{\mathrm{mem},k} = k_\mathrm{m} D_s^+ D_s^- \bar{\boldsymbol{X}}_k$, midpoint membrane stretch $\bar{J}_k$, force spreading, and fluid interpolation use $\bar{\boldsymbol{X}}$. The solute jump $[c]_k^n$ and F-actin concentration remain fixed during the mechanical iteration. The accepted F-actin field supplies the face drag coefficients, bulk force, and marker stress $\sigma_k^n = k_\sigma F_{\Gamma,k}^n$. Face values are extended from the accepted intracellular F-actin field and restricted by the accepted interior face mask. Membrane traces are retained by material marker index.

The water-permeation law in main-text Eq. (6) determines membrane motion relative to the fluid. The osmotic and elastic contributions use the old membrane normal $\boldsymbol{n}_k^n$ and are given, respectively, by

$$k_\mathrm{w} RT [c]_k^n \boldsymbol{n}_k^n, \qquad \frac{k_\mathrm{w}(\bar{\boldsymbol{F}}_{\mathrm{mem},k} \cdot \boldsymbol{n}_k^n)\boldsymbol{n}_k^n}{\bar{J}_k}.$$

The actin and adhesive contributions use the midpoint normal $\bar{\boldsymbol{n}}_k$ and are given, respectively, by

$$k_\mathrm{w} \sigma_k^n \bar{\boldsymbol{n}}_k, \qquad -k_\mathrm{w} k_\mathrm{ad}(\boldsymbol{V}_k^{n+1} \cdot \bar{\boldsymbol{n}}_k)\bar{\boldsymbol{n}}_k.$$

### S3. Moving-domain transport with Robin interface conditions

Solute, F-actin, and G-actin are advanced by the same Cartesian moving-domain method [16]. Both transport solves use $\boldsymbol{u}^{n+1}$, $\boldsymbol{X}^n$, $\boldsymbol{X}^{n+1}$, $\boldsymbol{n}^{n+1}$, $\Delta t$, and $\boldsymbol{V}^{n+1}$ from the accepted mechanical step, with transport geometry $\Gamma^{n+1}$. The solute solve holds this state fixed; after the solute concentrations are accepted, the actin solve receives the same state and holds it fixed. Fluid velocities at membrane markers use bilinear MAC interpolation $I_{\text{MAC}}^{n+1}$; concentration traces and normal derivatives use sharp local quadratic reconstruction. All time levels refer to one fixed computational frame; Section S4 specifies the translating-frame treatment. The method is stated for a generic field $\phi$ on one side $\Omega_\ell(t)$ of the membrane, where $\ell = i$ or $e$. The transport velocity is $\boldsymbol{v}$, diffusivity is $D$, reaction coefficient is $\kappa$, source is $f$, and the Robin coefficient is determined by the relative normal velocity of the membrane and transporting phase:

$$\partial_t\phi + \nabla\cdot(\boldsymbol{v}\phi) - D\Delta\phi + \kappa\phi = f \quad \text{in } \Omega_\ell(t), \qquad \partial_n\phi + \alpha\phi = g \quad \text{on } \Gamma(t), \qquad \alpha = \frac{(\boldsymbol{V}-\boldsymbol{v})\cdot\boldsymbol{n}}{D}. \tag{4}$$

Each one-sided problem is extended to the Cartesian box $B$, denoted by $\Omega$ in the main text, with continuity of the extended field and an unknown normal-derivative jump $\psi$:

$$\partial_t\phi + \nabla\cdot(\boldsymbol{v}\phi) - D\Delta\phi + \kappa\phi = f \quad \text{in } B, \qquad [\phi] = 0, \quad [\partial_n\phi] = \psi \quad \text{on } \Gamma(t). \tag{5}$$

The continuity condition $[\phi] = 0$ applies to the auxiliary extension for each one-sided problem. The physical solute concentrations retain separate interior and exterior traces and their osmotic jump. The interface density is fixed by the physical Robin condition,

$$\pm\tfrac{1}{2}\psi + \{\partial_n\phi\} + \alpha\,\{\phi\} = g \qquad \text{on } \Gamma(t). \tag{6}$$

At cell centers away from the membrane, backward Euler and centered conservative differences give

$$L_h^{n+1}\phi^{n+1} = D_t^-\phi^{n+1} + D_x^-\left(v^x A_x^+\phi^{n+1}\right) + D_y^-\left(v^y A_y^+\phi^{n+1}\right) - D\,\Delta_h\phi^{n+1} + \kappa\phi^{n+1} = f^{n+1}. \tag{7}$$

A stencil that crosses the membrane receives a correction derived from the one-sided smooth extensions. With $C_h$ the local quadratic correction function and $\chi$ the intracellular indicator,

$$L_h^{n+1}\phi^{n+1}_{i+\frac{1}{2},j+\frac{1}{2}} = f^{n+1}_{i+\frac{1}{2},j+\frac{1}{2}} + Q^{n+1}_{i+\frac{1}{2},j+\frac{1}{2}}, \qquad Q^{n+1} = L_h^{n+1}(\chi^{n+1}C_h^{n+1}) - \chi^{n+1}L_h^{n+1}C_h^{n+1}. \tag{8}$$

The correction is nonzero only at cut stencils, so the bulk coefficient matrix retains the same Cartesian structure throughout the domain. For the actin correction history, the old correction polynomial $C_h^n$ is evaluated at the updated marker position about its old center $\boldsymbol{X}_k^n$; the old nearest-marker map is retained for this evaluation in cells swept by the interface. The unknown interface density is solved with matrix-free GMRES; each matrix–vector product requires one bulk multigrid solve [16].

The solute Robin condition in main-text Eq. (7) prescribes transfer relative to the moving membrane. The relative normal velocity and transfer coefficients are

$$\begin{gathered} r_k^{n+1} = (I_{\text{MAC}}^{n+1}\boldsymbol{u}^{n+1} - \boldsymbol{V}^{n+1})_k \cdot \boldsymbol{n}_k^{n+1}, \\ A_k = k_c + \max(k_p(s_k), 0), \qquad B_k = -k_c + \min(k_p(s_k), 0). \end{gathered}$$

The cell-outward normal is used on both sides, and the solute-transfer expression is $A_k c_i + B_k c_e$. The exterior solve uses the accepted interior trace, retained by material marker index. The interior solve then uses the updated exterior trace. The corresponding Robin conditions, in solve order, are

$$\begin{gathered} -D_S\partial_n c_e^{n+1} + (r_k^{n+1} - B_k)c_{e,\Gamma,k}^{n+1} = A_k c_{i,\Gamma,k}^{n}, \\ -D_S\partial_n c_i^{n+1} + (r_k^{n+1} - A_k)c_{i,\Gamma,k}^{n+1} = B_k c_{e,\Gamma,k}^{n+1}. \end{gathered}$$

This exterior–interior pass gives $[c]_k^{n+1} = c_{i,\Gamma,k}^{n+1} - c_{e,\Gamma,k}^{n+1}$.

The F-actin bulk operator uses velocity $a\boldsymbol{u}^{n+1}$, diffusivity $D_F$, and sink $\gamma F^{n+1}$; the G-actin operator uses velocity $\boldsymbol{u}^{n+1}$, diffusivity $D_G$, and source $\gamma F^{n+1}$. Both bulk conversion terms vanish when $\gamma = 0$. The approximation to main-text Eq. (10) is

$$q_k^n = \frac{J_{\mathrm{poly}} g(s_k)}{G_{\Gamma,k}^n + \theta_{c,c}}, \qquad J_{\mathrm{actin},k}^{n+1} = q_k^n G_{\Gamma,k}^{n+1}.$$

The saturation denominator uses the accepted old G-actin trace retained by material marker index. The relative normal velocities give the coefficients

$$\beta_{F,k}^{n+1} = \frac{(\boldsymbol{V}^{n+1} - a I_{\mathrm{MAC}}^{n+1} \boldsymbol{u}^{n+1})_k \cdot \boldsymbol{n}_k^{n+1}}{D_F},$$
$$\beta_{G,k}^{n+1} = \frac{(\boldsymbol{V}^{n+1} - I_{\mathrm{MAC}}^{n+1} \boldsymbol{u}^{n+1})_k \cdot \boldsymbol{n}_k^{n+1}}{D_G}.$$

The boundary conditions are

$$\partial_n F^{n+1} + \beta_F^{n+1} F_\Gamma^{n+1} - \frac{q^n}{D_F} G_\Gamma^{n+1} = 0,$$
$$\partial_n G^{n+1} + \left(\beta_G^{n+1} + \frac{q^n}{D_G}\right) G_\Gamma^{n+1} = 0.$$

The choice $J_{\mathrm{poly}} = 0$ gives $q_k^n = 0$ and eliminates membrane polymerization.

The actin interface densities form one vector of length $2N_\Gamma$, where $N_\Gamma$ is the number of membrane markers. For each trial pair, the F-actin bulk solution supplies $\gamma F^{n+1}$ to the G-actin solve, and both interface residuals use the resulting G-actin trace. GMRES solves this joint affine system with paired polymerization fluxes. The final evaluation supplies the accepted F/G fields, traces, and correction histories.

### S4. Temporal coupling, resolution, and solver settings

At time $t^n$, one step first advances the semi-implicit fluid–membrane problem with the accepted solute jump and F-actin field. Solute and coupled F-actin/G-actin transport then use the fixed mechanical state described in Section S3. The updated jump $[c]^{n+1}$ and F-actin field provide osmotic feedback, drag coefficients, bulk force, and membrane stress for the next mechanical step; updated G-actin provides the saturation denominator for the next actin step. Backward Euler transport and lagged chemical feedback make the complete coupled method first order in time [10]. The bulk spatial operators are second order, and the moving-domain correction has second-order local spatial consistency [16].

**Supplementary Table S5. Numerical resolutions and sampling.**

| Calculation | Domain, grid, membrane discretization, and simulation interval |
| --- | --- |
| Figure 2A–C parameter sweeps | $100 \times 100$ μm$^2$ translating box; $128^2$ cells; spacing 0.781 μm; 120 membrane markers; nominal 5.00 s time step. Velocity is reported at 450 s from the 425–475 s centroid displacement. Selected Figure 2C cases use 2.50 or 1.25 s controls when the 5.00 s run does not supply the complete reporting interval. |
| Figure 2D conservation | Same spatial discretization as Figure 2A–C; 5.00 s time step; simulation through $1.00 \times 10^4$ s; fields saved every 50.0 s. |
| Figure 2E | Same $100 \times 100$ μm$^2$, $128^2$, 120-marker discretization; field shown at $2.00 \times 10^3$ s; archived calculation extends to $4.00 \times 10^3$ s. |

| Calculation | Domain, grid, membrane discretization, and simulation interval |
|---|---|
| Figure 2F | Same $100 \times 100$ μm$^2$, $128^2$, 120-marker discretization; field shown at $5.00 \times 10^2$ s. |
| Pump-induced calculations | $80 \times 80$ μm$^2$ translating box; $128^2$ grid; spacing 0.625 μm; 300 membrane markers; 5.00 s time step; histories and terminal fields through $2.00 \times 10^3$ s. |
| Coupled calculations (Fig. 4A–K) | $100 \times 100$ μm$^2$ box with horizontal recentering of membrane markers; $128^2$ cells; 120 membrane markers; $\Delta t = 5$ s. Area, polarity, and displacement are sampled every 5 s; fields and reconstructed substrate force every 50 s. Maps and velocity profiles use 2000 s. |
| Actin-driven spatial refinement | $128^2/120$, $256^2/240$, and $512^2/480$ grid-cell/marker configurations; $\Delta t = 1.25$ s; 0–2000 s. |
| Actin-driven temporal refinement | $128^2$ cells and 120 markers; $\Delta t = 5$, 2.5, and 1.25 s; 0–2000 s. |
| Central-profile matched refinements | $256^2/240$, $256^2/480$, $512^2/480$, and $512^2/960$ at 1.25 s; additional $512^2/480$ calculation at 0.625 s. All use profile K12 and the four matched conditions. |
| Area and force profile checks | All five pump profiles at $256^2$ cells, 480 markers, and 1.25 s steps, compared with the main resolution over 0–2000 s. |

After each accepted fluid–membrane update, the coupled calculations shift all membrane markers in the periodic $x$-direction so that their mean $x$-coordinate equals the box midpoint. The MAC velocity and pressure arrays retain the values returned by the mechanical solve. Both transport solves use that MAC velocity field and the old and recentered new marker positions. The Robin conditions use normals evaluated on the recentered membrane and membrane velocities given by the marker displacements from the old to the recentered positions divided by $\Delta t$. To obtain the laboratory-frame area-centroid position, we add the accumulated horizontal shift to the polygon area centroid in the computational box. The velocity profiles in Fig. 4G,H report laboratory cytosolic and F-actin velocities relative to the laboratory area-centroid velocity on the exact centroid centerline; centroid velocity is obtained from the nominal centered 200 s fitting window, truncated at the final time. At 2000 s, the three laboratory centroid positions at 1900, 1950, and 2000 s give

$$V_x = \frac{x_{\mathrm{c}}^{\mathrm{lab}}(2000) - x_{\mathrm{c}}^{\mathrm{lab}}(1900)}{100\ \mathrm{s}}.$$

MAC face velocities are averaged to cell centers before evaluation on the centroid line, and $V_x$ is subtracted from both cytosolic and F-actin $x$-velocities.

The nonlinear membrane residual tolerance is $10^{-10}$. The interface transport solves use relative GMRES tolerances of $10^{-9}$–$10^{-10}$, and multigrid solves use a tolerance of $10^{-12}$. For the parameter sweeps (Fig. 2A,B), a parameter location is retained when all three $k_{\mathrm{ad}}$ branches contain the complete 425–475 s interval; C uses each available branch independently. The zero-polymerization point is omitted because one branch does not reach the reporting interval.

### Spatial, temporal, and membrane refinement

Refinement tests use the physical parameters of the coupled calculations and the central pump profile K12 (Tables S1, S2, and S4). Physical parameters and initial conditions are held fixed within each simulation condition; numerical configurations are listed in Table S5.

At common saved times $t_i$, the coarser and finer histories $q_c$ and $q_f$ give

$$D_\infty(q) = \max_i |q_c(t_i) - q_f(t_i)|,$$
$$D_{RMS}(q) = \left(\frac{1}{n}\sum_i [q_c(t_i) - q_f(t_i)]^2\right)^{1/2},$$
$$\delta_\infty(q) = 100\,\frac{D_\infty(q)}{\max_i |q_f(t_i)|}.$$

Here $n$ is the number of common samples, and $\delta_\infty(q)$ is expressed as a percentage over the same interval. The observable $q$ is actin-driven displacement or excess displacement $\Delta d = d_{AW} - d_A - d_W + d_0$.

Joint grid/marker refinement of actin-driven migration reduces both the maximum and RMS displacement differences, with observed rates of 1.54 and 1.48, respectively (Fig. S1A). Temporal refinement gives corresponding rates of 1.03 and 1.00, consistent with the first-order coupled method (Fig. S1B). The rates are calculated as $p = \log_2(D_{01}/D_{12})$, where $D_{01}$ and $D_{12}$ compare successive resolution levels using the same norm. An additional time-step halving on the finest grid changes actin-driven displacement by at most 0.002347 µm, or 0.01587% of the refined maximum; this actin-driven calculation meets the numerical quality criteria over the full reporting interval.

The quality criteria require absolute relative changes in total actin and total solute of at most 0.1%, neighboring membrane-marker distances between $0.5h$ and $2.5h$, and a largest-to-smallest distance ratio of at most 4, where $h = (100/N)$ µm for an $N^2$ grid. The membrane must remain non-self-intersecting, saved intracellular F-actin and G-actin must be nonnegative within numerical tolerance, and laboratory-frame positions must be verified. Complete-interval comparisons require every common sample. Two coarse baseline time-refinement runs that stopped at 50 and 370 s under the solver stagnation criterion are excluded from the excess-displacement comparisons. The three-level rates above concern actin-driven displacement; the two-level matched comparisons below measure the sensitivity of the interaction contrast to numerical resolution.

Joint grid/marker, time-step, and marker-only refinements give consistent excess-displacement histories over the common interval in which all matched conditions satisfy the numerical quality criteria (Table S6; Fig. S1C). This interval precedes the first membrane-spacing departure at 605 s among the standard refined combined configurations. Across all four conditions and the refined configurations, the largest actin and solute conservation changes over this interval are 0.0222% and 0.00674%, respectively. The grid comparison with denser membrane markers meets all criteria through 1000 s; its maximum excess-displacement difference over this interval is 0.008647 µm, or 0.5372%. The refined displacement histories retain the positive interaction between active solute pumps and actin turnover (Fig. S1D).

**Table S6:** Refinement sensitivity of excess displacement for profile K12. Joint grid/marker and marker-only comparisons use 1.25 s steps. $N^2/M$ denotes grid cells/membrane markers. The two intervals contain 121 and 401 common samples.

| Comparison | $D_\infty$ (µm) 0–600 s | $\delta_\infty$ (%) 0–600 s | $\delta_\infty$ (%) 0–2000 s |
|---|---|---|---|
| $256^2/240$ vs. $512^2/480$ | 0.003650 | 0.4875 | 0.3467 |
| $256^2/480$ vs. $512^2/960$ | 0.005275 | 0.7039 | 0.4805 |
| Time step: 1.25 vs. 0.625 s on $512^2/480$ | 0.002388 | 0.3180 | 0.2744 |
| Markers: 240 vs. 480 on $256^2$ cells | 0.002467 | 0.3269 | 0.8774 |
| Markers: 480 vs. 960 on $512^2$ cells | 0.000841 | 0.1123 | 0.1937 |

At the main resolution, the central combined calculation first exceeds the membrane-spacing criteria at 595 s. By 2000 s, its actin and solute conservation errors reach 0.925% and 0.157%, and the pump-induced control has a 0.266% actin error. On the finest grid with denser markers, the earliest membrane-spacing

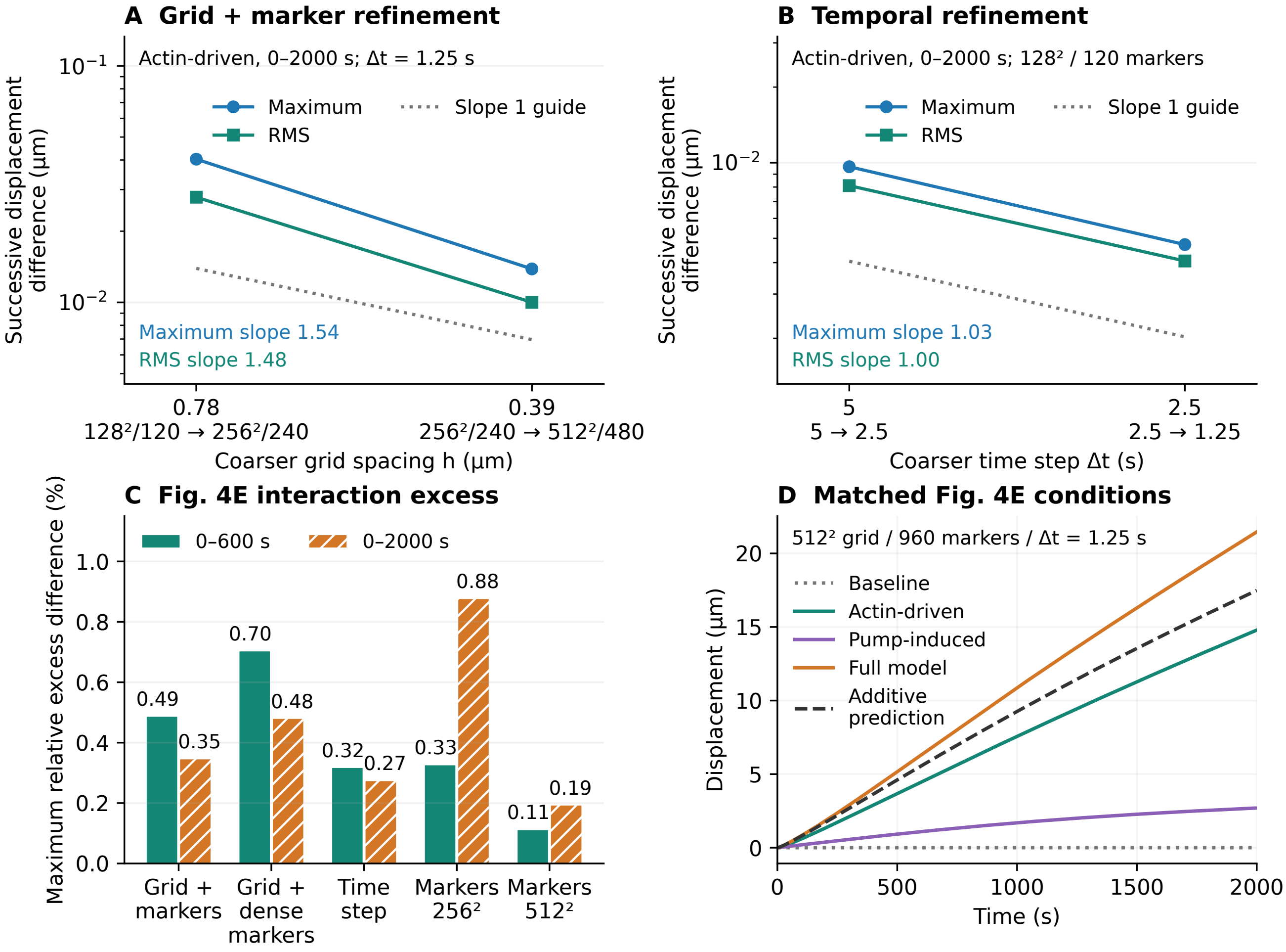


**Figure S1: Spatial, temporal, and membrane refinement of migration.** **(A,B)** Successive maximum and RMS differences in actin-driven displacement under joint grid/marker refinement and time-step refinement over 0–2000 s. **(C)** Normalized maximum differences in excess displacement for profile K12. Solid and hatched bars use 0–600 s and 0–2000 s; the longer interval includes departures from the numerical quality criteria. **(D)** Matched displacements, including the combined condition labeled Full model, and their baseline-corrected sum on $512^2$ cells with 960 markers and 1.25 s steps. All four conditions meet the criteria through 1000 s. Numerical configurations and quantitative comparisons are given in Tables S5–S7.

departure across the four conditions occurs in the actin-driven calculation at 1191.25 s; the combined spacing departure occurs at 1351.25 s, while its maximum actin-conservation error over the full reporting interval is 0.05725%. Excess displacement at 2000 s ranges from 3.95872 to 3.99376 μm across the refined configurations. The comparisons in Table S7 include these late departures. Individual displacements are reported alongside the interaction contrast because subtraction can partially cancel their numerical differences.

**Table S7:** Main-resolution and refined central-profile displacement comparisons over 0–2000 s. Main and refined configurations are $128^2/120$ at 5 s and $512^2/960$ at 1.25 s. Endpoint values and $D_\infty$ are in μm. Relative differences use the maximum absolute refined value over the same interval.

| Quantity | Main endpoint | Refined endpoint | $D_\infty$ | $\delta_\infty$ (%) |
|---|---|---|---|---|
| Actin-driven | 14.72066 | 14.78894 | 0.06828 | 0.4617 |
| Pump-induced | 2.73616 | 2.69754 | 0.05878 | 2.1792 |
| Combined | 21.31501 | 21.46115 | 0.14613 | 0.6809 |
| Excess $\Delta d$ | 3.85819 | 3.97467 | 0.11647 | 2.9304 |

Joint refinement of the combined pump profiles changes relative-area and signed substrate-force histories by at most 3.89% and 2.65%, respectively, over the full reporting interval, using the configurations in Table S5. Here $q = A/A_0$ or $T_x$ in the definition of $\delta_\infty(q)$. For the widest profile, H140, normalization of the endpoint area difference by the refined area increase $A(2000)/A_0 - 1$ gives 5.13%. These refined profile checks satisfy the conservation criterion, while membrane-spacing departures remain at late times.

### S5. Conservation, polarity, and matched-control diagnostics

Total actin and total solute are reconstructed from the saved physical fields. Fully intracellular or extracellular cells contribute their full area. Cells intersected by the membrane use geometric cell fractions and one-sided reconstructed concentrations integrated over the appropriate polygon. Total actin is $M_A = M_F + M_G$ over the intracellular domain. Total solute is $M_S = M_{\text{inside}} + M_{\text{outside}}$ over the full computational box. Relative conservation error is reported as $100[M(t) - M(0)]/M(0)$. These diagnostics quantify material conservation and are distinct from spatial or temporal convergence tests.

The Figure 2D actin tests extend to 10,000 s. Using fields saved every 50 s, the maximum absolute relative conservation errors are 0.0197%, 0.0101%, and 0.00551% for $k_{\text{ad}} = 8.00 \times 10^{-4}$, $1.00 \times 10^{-3}$, and $1.25 \times 10^{-3}$ Pa s/μm, respectively. The corresponding total-solute changes remain below 0.015% in these passive-pump calculations. In the pump-induced conservation test, the relative error in total solute remains below 0.01% through 2000 s for the equal-rear, twofold-wider-rear, and threefold-wider-rear profiles.

The coupled polarity diagnostic uses equal-weight intracellular grid samples in the front and rear regions. For either F-actin or G-actin, $P = (\bar{c}_{\text{front}} - \bar{c}_{\text{rear}})/(|\bar{c}_{\text{front}}| + |\bar{c}_{\text{rear}}|)$, with the instantaneous area centroid defining the split. This quantity measures concentration asymmetry and is distinct from the area-weighted conservation calculation.

The additional geometric diagnostic is $A(t)/A_0$, where $A(t)$ is the enclosed polygonal area and $A_0 = A(0)$. The signed horizontal force exerted on the substrate per unit out-of-plane thickness and the reconstructed laboratory-frame network flux are

$$T_x(t) = \int_{\Omega_i(t)} \eta_{\text{st}} \theta_n v_{n,x} \, \mathrm{d}A,$$

$$\theta_n \boldsymbol{v}_n = a\theta_n \boldsymbol{u}_{\text{lab}} - D_n \nabla \theta_n.$$

The force is reported in pN/μm of thickness and evaluated using the intracellular fractional-cell quadrature described above. Negative $T_x$ denotes rearward force on the substrate. This diagnostic includes the distributed actin–substrate drag; the membrane normal-resistance term governed by $k_{\text{ad}}$ contributes separately to the mechanical balance. Force curves connect the saved field times listed in Table S5.

The four matched controls share the same initial actin pools and passive material properties; membrane polymerization, bulk depolymerization, and active solute pumping are switched as specified in Table S3. All four conditions retain F-actin network stress and mechanical drag. The baseline-corrected additive displacement is $d_{\text{add}} = d_A + d_W - d_0$, and excess displacement is $\Delta d = d_{AW} - d_{\text{add}}$. Positive $\Delta d$ identifies migration beyond $d_A + d_W - d_0$ and quantifies the net nonlinear interaction between active solute pumping and actin dynamics. The contrast includes coupled changes in actin transport, F-actin network stress, drag, fluid flow, and cell geometry.

The coupled calculations share a saved endpoint of 2000 s across the twelve matched cases. Field maps, polarity, displacement, excess displacement, relative area, substrate force, and centerline velocity profiles use this common interval. Run-status diagnostics record conservation errors, minimum actin concentrations, membrane-marker spacing, boundary clearance, and frame-centering error. Saved actin fields retain any computed negative values so that the diagnostics expose loss of positivity.
